# Comparing neurophysiological correlates of speech-in-noise perception across MEG, EEG and ear-EEG

**DOI:** 10.64898/2026.07.31.741889

**Authors:** Till Habersetzer, Andreas Radeloff, Bernd T. Meyer

**Affiliations:** Communication Acoustics, Department of Medical Physics and Acoustics and Cluster of Excellence “Hearing4all”, Carl von Ossietzky Universität Oldenburg, Oldenburg, Germany; Division of Otolaryngology, Head and Neck Surgery, Carl von Ossietzky Universität Oldenburg, Oldenburg, Germany

## Abstract

Neural speech tracking is a powerful phenomenon in which brain signals synchronize with the time-varying features of continuous speech. This study benchmarks speech-in-noise tracking performance and the feasibility of objective speech reception threshold (SRT_neuro_) estimation across three sensor modalities: magnetoencephalography (MEG), scalp-electroencephalography (EEG), and ear-EEG. While MEG and scalp-EEG serve as reference measurements providing full-head coverage, ear-EEG offers a more unobtrusive and portable alternative. Twenty-one young, normal-hearing adults participated in simultaneous MEG and 76-channel EEG recordings, which incorporated 16 integrated around-ear electrodes (ear-EEG). During the recordings, participants listened to continuous audiobooks and matrix sentences at fixed intelligibility levels corresponding to individually varied signal-to-noise ratios (SNRs). Neural tracking was quantified via envelope reconstruction using a linear decoder, and psychometric functions were fitted to the data to derive SRT_neuro_ as the midpoint. We observed consistent envelope tracking across all three modalities, suggesting shared underlying neural processes, with reconstruction accuracies increasing alongside SNR. MEG yielded the highest signal quality, with reconstruction accuracies approximately 1.5 times higher than scalp-EEG and 3 times higher than ear-EEG. Although SRT_neuro_ was derived within a 5 dB margin of behavioral thresholds for all participants, these neurophysiological estimates did not significantly correlate with individual behavioral SRTs. Furthermore, ear-EEG proved less reliable and exhibited a slight bias. These results demonstrate that MEG, scalp-EEG, and ear-EEG are all viable for capturing robust neural envelope tracking. However, the acoustic envelope alone may be insufficient as a standalone predictor for diagnostics due to high inter-subject variability.

**Significance Statement:** When listening to speech, the brain tracks the rhythm of the signal, specifically the temporal envelope. These neural responses provide insights into speech-in-noise perception and offer a potential objective measure of listening ability. This study is the first to benchmark speech-in-noise tracking across high-resolution MEG, affordable scalp-EEG, and emerging ear-EEG using individualized signal-to-noise ratios derived from psychometric functions. Our comparison reveals a clear performance hierarchy, proving that even ear-centered configurations can capture robust neural signals across varying acoustic conditions relative to high-fidelity references. While high inter-subject variability currently limits the diagnostic utility of standalone envelope tracking, characterizing these trade-offs across different sensor modalities establishes a foundation for advanced, individualized neural hearing assessments.

## Introduction

Neural tracking of speech refers to the synchronization of neural activity with the time-varying features of continuous speech (Wöstmann et al. 2017; Brodbeck and Simon 2020). These features span a hierarchy ranging from low-level acoustic properties, such as amplitude and spectrum, to higher-level linguistic structures, including phonemes, words, and phrases (Brodbeck et al. 2018b; Gillis et al. 2023). This phenomenon facilitates the use of ecologically valid stimuli, such as continuous natural speech, and enables various practical applications. A prominent example is Auditory Attention Decoding (AAD), which aims to develop closed-loop, neuro-steered hearing aids (Geirnaert et al. 2021). By analyzing brain activity, these systems would identify and selectively enhance the voice of the speaker to whom the user is attending.

Furthermore, neural tracking is increasingly utilized as an objective tool for assessing the auditory pathway, particularly in its relationship to behavioral measures such as speech comprehension (Gillis et al. 2022). Understanding speech in noisy environments is a major challenge that impacts quality of life across the lifespan (WHO et al. 2021). This ability is traditionally assessed via the behavioral speech reception threshold (SRT), the signal-to-noise ratio (SNR) at which 50 % of speech is understood. While behavioral tests require active participant feedback, objective neural measures could provide access to non-responsive populations (e.g., infants or unconscious patients), facilitate hearing aid optimization through improved noise reduction, and enable long-term monitoring of speech intelligibility. Previous studies have demonstrated the potential for predicting SRTs using electroencephalography (EEG) (Vanthornhout et al. 2018; Lesenfants et al. 2019; Muncke et al. 2022; Borges et al. 2025b; Deoisres et al. 2025) and specifically ear-EEG (Borges et al. 2025c), yet significant variability remains, complicating clinical application.

Investigating time-varying neural responses to complex stimuli like speech requires methods with high temporal resolution, such as EEG or magnetoencephalography (MEG) (Crosse et al. 2016). These non-invasive techniques provide millisecond-scale resolution and directly measure neural activity. Both methods provide whole-head coverage, though MEG is less susceptible to volume conduction than EEG. Since the skull, scalp and other tissues are nearly transparent to magnetic fields, MEG signals exhibit significantly less spatial smearing and distortion. This allows for a more direct interpretation of sensor topography and higher-precision source imaging, making MEG a superior tool for resolving large-scale brain dynamics (Baillet 2017; Gross 2019). However, MEG entails substantial capital and operating costs comparable to an MRI scanner, and measurements are restricted to magnetically shielded rooms. In contrast, EEG remains affordable, portable, and easily maintainable (Nicolas-Alonso and Gomez-Gil 2012). Simultaneous MEG and EEG recordings are often recommended because the two measures are complementary (Malmivuo 2012). They differ in their sensitivity to neural current orientations. EEG captures both radial and tangential components, while MEG is predominantly sensitive to tangential currents (Silva 2013). Combining these modalities provides a more comprehensive representation of neural generators and improves source localization accuracy. However, such combined recordings entail technical and practical challenges. In contrast to full-head methods, ear-EEG offers an unobtrusive, more portable alternative for long-term monitoring in everyday environments (Looney et al. 2012; Debener et al. 2012; Debener et al. 2015). Ear-EEG is particularly effective for targeting auditory processes, as it demonstrates high sensitivity to sources within the temporal lobe (Meiser et al. 2020).

This study benchmarks the performance of neural tracking and SRT prediction across three modalities: MEG, scalp-EEG, and ear-EEG. To our knowledge, this is the first study to evaluate these metrics across all three modalities within a unified experimental framework. We hypothesized that performance would be most robust for MEG, followed closely by scalp-EEG, while ear-EEG would remain a viable and feasible alternative. All three modalities were recorded simultaneously, with ear-EEG data captured via around-ear electrodes integrated into the a EEG-cap. In this configuration, high-density MEG and scalp-EEG serve as complementary, full-head reference methods, allowing us to evaluate the performance of ear-EEG as a more practical and unobtrusive alternative. Consistent with established research, we quantified neural tracking using reconstruction of the envelope via a linear decoder as the speech amplitude envelope is robustly linked to speech intelligibility (Luo and Poeppel 2007; Peelle and Davis 2012; Pasley et al. 2012), and linear models remain a standard method for assessing neural tracking (Crosse et al. 2021). To predict SRTs, decoders were trained on continuous audiobooks and applied to matrix sentences across varying SNRs (Vanthornhout et al. 2018). We quantitatively compared the three modalities based on reconstruction performance, SRT prediction feasibility, and inter-subject variability to assess their practical utility.

## Materials and Methods

### Participants

Twenty-one native German speakers (12 females, 9 males; age: 27 *±* 4 years) were recruited from the University Campus. All participants reported no history of neurological disorders and had normal or corrected-to-normal vision. Prior to participation, subjects were screened for normal hearing using a pure-tone audiogram. Normal hearing was defined as a four-frequency (0.5, 1, 2, 4 kHz) pure-tone average (4fPTA) below 20 dB HL for the better ear (Humes 2019). All participants met this criterion. However, two individuals showed slightly increased 4fPTAs in their other (non-better) ear: sub-01 (left ear: 20 dB HL) and sub-19 (right ear: 21.25 dB HL). Both participants performed normally in subsequent speech audiometry and reported no subjective hearing loss. As handedness was not formally assessed, no related exclusion criteria were applied. Participants provided written informed consent regarding the study procedure and the open sharing of their anonymized data. They were compensated for their time. The study was approved by the University Ethics Committee (Ref: EK/2021/181) and conducted in compliance with the Declaration of Helsinki.

### Experimental design

The study followed a three-session design per participant: one screening session (ses-00) and two recording sessions (ses-01, ses-02). Data collection for the entire cohort spanned a period of four months.

#### Session 0: Screening and preparation

The first session (approx. 2 h) served to screen participants and familiarize them with the MEG environment. Following the standard pure-tone audiogram in a sound-attenuated booth, participants were positioned inside the MEG to assess comfort, head size compatibility, and potential artifacts for MEG compatibility. Subsequently, two runs of speech audiometry were conducted inside the MEG, along with the measurement of transient hearing thresholds for click and up-chirp stimuli. This session also included sizing for the EEG cap and custom earmolds. Structural MRI scans were acquired either at the end of this session or postponed until after the following recording sessions depending on scanner availability. Since the transient click/chirp stimuli and MRI scans are not within the scope of the research questions posed in this paper, they are not analyzed in the present study.

#### Sessions 1 & 2: MEG/EEG recording

The subsequent two sessions (approx. 3 h to 4 h each) focused on simultaneous MEG and EEG data acquisition. Each session began with participant preparation (*≈* 1 h), followed by data collection inside the magnetically shielded room (*≈* 1.5 h to 2 h). Once positioned in the MEG, participants completed two runs of speech audiometry to determine their individual behavioral speech reception threshold (SRT_beh_). These thresholds were used to calibrate the SNRs for the subsequent experimental blocks.

### Paradigm and stimuli

#### Behavioral audiogram and speech audiometry (OLSA)

- **Audiogram:** Pure-tone air conduction thresholds were measured for the left and right ears separately. Testing covered frequencies 0.125, 0.25, 0.5, 0.75, 1, 1.5, 2, 3, 4, 6 and 8 kHz. All measurements were conducted in a sound-attenuated booth using an Oscilla A50 audiometer (Oscilla, Aarhus, Denmark).
- **OLSA:** Speech intelligibility was assessed using the German Oldenburg Sentence Test (OLSA) (Wagener et al. 1999; Wagener and Brand 2005). We employed an open-access version featuring synthesized female speech from the corpus described by Nuesse et al. (2019), which has been validated to yield thresholds comparable to natural speech. The test material is based on a matrix of 50 words (10 alternatives for each of the five syntactic positions). Each sentence follows the fixed structure: *Name–verb–numeral–adjective–object*. Sentences are generated as pseudo-randomized paths through this matrix and arranged into lists of 20, which are phonetically balanced to yield equivalent intelligibility scores. Sentences were presented in continuous, stationary speech-shaped noise. This noise was generated from the speech material itself, ensuring it shared the same long-term frequency spectrum as the sum of all sentences. The background noise level was fixed at 65 dB SPL. Participants were instructed to repeat the words they understood after each sentence. An adaptive procedure adjusted the speech presentation level to determine the SRT_beh_, defined as the SNR yielding 50 % correct word recognition. OLSA measurements were conducted inside the magnetically shielded room underneath the MEG helmet using insert earphones and the intercom of the MEG. Testing occurred at the beginning of each session, directly before MEG/EEG acquisition. In total, six measurements were obtained per subject (two per session). The SRT_beh_ determined from the second measurement of the first recording session (ses-01) served as the reference value to calculate the SNRs for the target intelligibility levels in all subsequent experiment blocks (ses-01 and ses-02). Therefore, prior to data collection, participants completed three training lists split across two days to familiarize themselves with the procedure and stabilize their performance. This aligns with established guidelines, which typically recommend one to two training lists (Wagener et al. 1999; Nuesse et al. 2019). The experiments were controlled using a custom Python implementation of the OLSA. The software and stimulus material will be made openly available upon publication to ensure full replicability.

#### Experimental tasks

The main experiment consisted of three listening tasks presented during simultaneous MEG/EEG recordings. This paper focuses on two of these tasks (excluding transient task): the Oldenburg Sentence Test and audiobooks. The task order was identical for every session: two blocks of OLSA, followed by two blocks of audiobooks and one final OLSA block.

- **Audiobooks:** Stimuli consisted of two German public-domain short stories: “The Tell-Tale Heart” (E.A. Poe) and “The Stolen Bacillus” (H.G. Wells). The audio was generated using a neural text-to-speech engine (ElevenLabs), utilizing a custom voice model based on a female speaker from the research group. The clean speech recordings were presented at a long-term average RMS level of 65 dB SPL. Each story lasted approximately 16 min and was divided into two blocks of equal duration. To ensure vigilance, participants answered three comprehension questions after each block. The presentation order was fixed, with “The Tell-Tale Heart” presented in the first session and “The Stolen Bacillus” in the second. Participants were ask to fixate a cross on the screen during the task.
- **OLSA:** The OLSA task was divided into three blocks containing a total of 540 sentences per session (200 sentences in each of the first two blocks, and 140 in the last). Sentences were derived from standard OLSA lists and mapped to six fixed intelligibility levels: 0 %, 20 %, 40 %, 50 %, 80 % and 100 %. The 0 % condition was included as a supplementary level. To balance the increase in total experiment duration, it was restricted to 40 sentences (two lists), while the five non-zero levels were assigned 100 sentences (five lists) each, resulting in a total of 27 lists per session. The order of list/SNR pairings was randomized, but sentences were always presented in concatenated groups of 20 (one full list) at a constant SNR. These 27 list combinations were distributed across the three blocks (10 lists in Block 1, 10 in Block 2, and 7 in Block 3). To avoid frustration and ensure awareness of playback onset, each block began with a list at an intelligibility level *≥* 50 %. To monitor attention, continuous playback was interrupted by response trials. A response was mandatory after each completed list of 20 sentences. Additionally, within each list, there was a 60 % probability of an intermediate response. The position of this intermediate response was determined by drawing from a normal distribution (*µ* = 10, *σ* = 4), restricted to sentence indices between 6 and 14 to prevent interruptions near the list boundaries. Participants responded via a button box (RESPONSEPixx/MRI, VPIXX, Saint-Bruno-de-Montarville, Canada) using a 2-alternative forced-choice paradigm. Two words were displayed: the correct target word and one incorrect alternative. A response was required within 4 s. Otherwise, the trial was marked as incorrect. Visual feedback (green/red) was provided immediately after the decision. Between response trials, sentences were concatenated with an inter-stimulus interval (ISI) jittered uniformly between 0.8 s and 1.2 s. Continuous OLSA background noise was fixed at 65 dB SPL. The onset of the first sentence in each continuous trial was jittered between 1.0 s and 1.5 s. SNRs were varied by adjusting the speech level. The mapping between target intelligibility and SNR was based on the individual psychometric function (Brand and Kollmeier 2002):

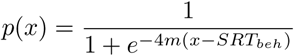

Here, *p*(*x*) is the probability of correct recognition at SNR *x*. The speech reception threshold (*SRT_beh_*) was estimated individually for each subject inside the MEG prior to the main experiment. The slope *m* was fixed at 0.13 for all subjects (Nuesse et al. 2019). While intermediate intelligibilities were mapped to SNRs using the fitted function (20 %: SRT_beh_ - 2.7 dB, 40 %: SRT_beh_ - 0.8 dB, 50 %: SRT_beh_, 80 %: SRT_beh_ + 2.7 dB), the anchor points were fixed and added to capture floor and ceiling effects: 0 % intelligibility was mapped to *−*40 dB, and 100 % intelligibility was mapped to 0 dB. Prior to data collection, participants completed a training run (two lists, 40 sentences) to familiarize themselves with the task. Participants were ask to fixate a fixation cross on the screen during the task.

#### Subject-specific exclusions and protocol deviations

The final experimental design (including 3 sessions, continuous Head Position Tracking (cHPI), and individualized SRT_beh_ values) was fully implemented from sub-02 onwards. Data from sub-00 (pilot) and sub-01 were included in the analysis despite the following protocol deviations. Subject sub-00 completed a preliminary version of the paradigm using fixed SNRs derived from a psychometric function (SRT_beh_: *−*9.6 dB, slope: 0.13), rather than individualized SNRs. Additionally, cHPI was turned off and standard pre-auricular points were used for coregistration instead of the earmold procedure employed for all subsequent subjects. Furthermore, sub-00 included only 5 of the 6 final SNRs, omitting the *−*40 dB condition and a clean speech condition (no noise) was used instead of the 0 dB SNR condition. Subject sub-01 adopted the final paradigm (individualized SRT_beh_ value, cHPI on, earmold co-registration) but utilized a lowest SNR of *−*20 dB instead of the *−*40 dB condition used for all subsequent participants. Both, sub-00 and sub-01 followed a two-session design, whereas the final design employed from sub-02 onwards consisted of three sessions. Consequently, for the first two subjects, the screening was integrated into the first MEG/EEG recording session, no behavioral OLSA data were collected for screening, and training consisted of 1 list instead of 3 OLSA lists.

**Table 1:** Protocol deviations for early subjects compared to the final experimental design.

| Feature | sub-00 (Pilot) | sub-01 | sub-02 Onwards |
| --- | --- | --- | --- |
| SRT<sub>beh</sub> Type | Fixed (−9.6 dB) | Individualized | Individualized |
| Highest SNR Cond. | Clean Speech | 0 dB | 0 dB |
| Lowest SNR Cond. | N/A (5 conditions) | −20 dB | −40 dB |
| Total Sessions | 2 | 2 | 3 |
| Training Lists | 1 | 1 | 3 |
| cHPI | No | Yes | Yes |
| Co-registration | Pre-auricular | Earmold | Earmold |
| Screening | Mixed in Session 1 | Mixed in Session 1 | Separate Session |

### Data acquisition

#### MEG and EEG

Neurophysiological data were recorded using a combined MEG/EEG setup in a magnetically shielded room (Vacuumschmelze, Hanau, Germany). MEG measurements were conducted using an Elekta Neuromag Triux System (Elekta Oy, Helsinki, Finland) equipped with 306 channels. The sensors are arranged in 102 triplets, each comprising one magnetometer and two orthogonal planar gradiometers. Measurements were performed with subjects seated in an upright position (68*^◦^*). Simultaneous EEG was recorded using a custom-made MEG compatible 76-channel cap (Easycap, Wörthsee, Germany) connected directly to the MEG acquisition system. The layout followed the standard Elekta 64-channel arrangement, augmented with 12 additional electrodes placed in the vicinity of the ears (6 per side) to improve coverage of auditory areas (see Figure 1). During recording, the reference electrode was placed on the right side of the nose, and the ground electrode was placed on the forehead. Impedances were maintained below 5 kΩ whenever possible. Additionally, three bipolar channels were recorded to monitor physiological artifacts: one electrocardiogram (ECG) channel and two electrooculography (EOG) channels (vertical: VEOG, horizontal: HEOG). Head positions were continuously monitored using five HPI coils attached to the EEG cap in custom-made holders. Prior to the measurement, we digitized the positions of the HPI coils, the EEG electrodes, three anatomical landmarks (nasion, left/right pre-auricular points), and *>* 200 head-shape points using a Polhemus Fastrak device (Polhemus, Colchester, VT, USA). All signals were sampled at 1000 Hz with an online band-pass filter of 0.1 Hz to 330 Hz. The MEG signals were recorded without internal active shielding. To account for the participants’ tendency to slouch during recordings, they were instructed to reposition their heads against the top of the MEG helmet before each recorded task. This ensured consistent head positioning across the session.

**Figure 1:**
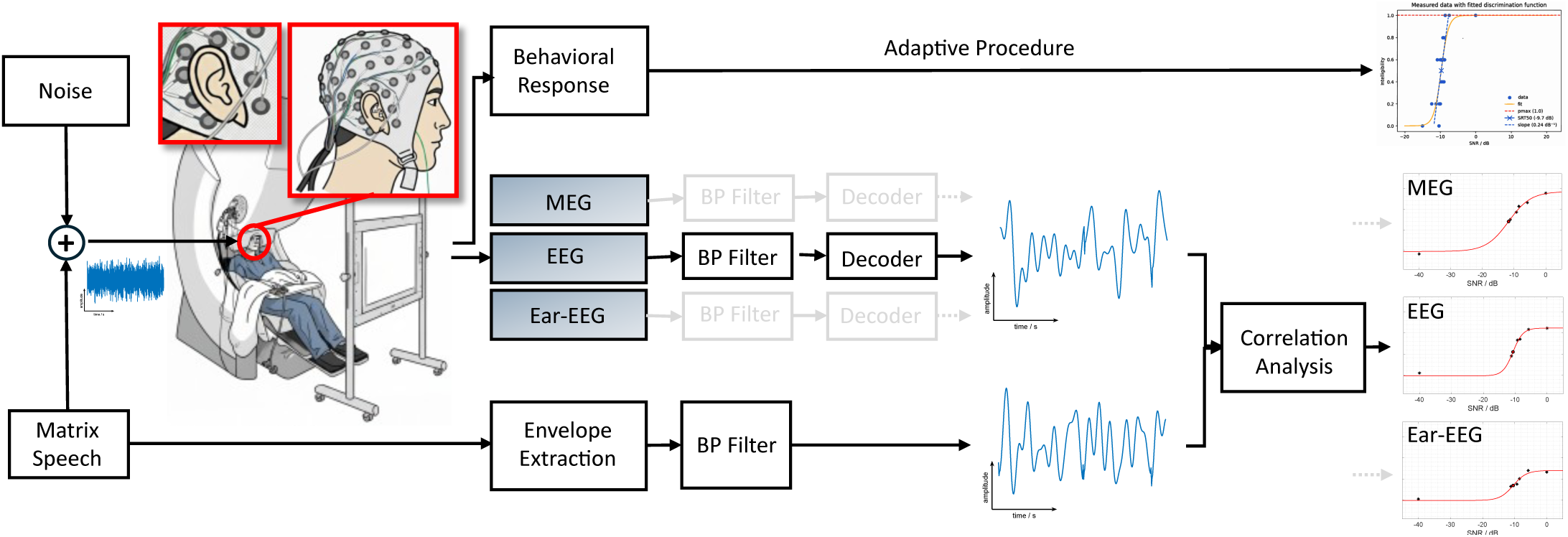
Overview of the experimental setup and analysis pipeline. Participants were seated in the MEG scanner while wearing a high-density EEG-cap featuring a subset of electrodes positioned around the ears (termed ear-EEG). MEG and EEG data were acquired simultaneously. For the analysis, we distinguish between three sensor modalities: the ear-EEG subset, the full EEG-cap (including all ear channels), and the MEG data. Acoustic stimulation was delivered via insert earphones and consisted of OLSA sentences presented at varying SNRs. Prior to the MEG/EEG recording, the SRT_beh_ was determined via an adaptive speech audiometry procedure inside the MEG using the same setup. For the data analysis, recordings from each modality were bandpass filtered and processed by a linear decoder trained on clean-speech audiobooks to reconstruct the speech envelope. Simultaneously, the true speech envelope was extracted from the audio and filtered. Neural envelope tracking was quantified by calculating the correlation between the reconstructed and true envelopes. These values, determined across different SNRs, were fitted with a sigmoid psychometric curve to derive the SRT_neuro_, which served as the neurophysiological analog to the SRT_beh_.

#### Hardware and software: Acoustic stimulation, calibration, and trigger details

Audio stimuli were delivered diotically using insert earphones (CareFusion Model TIP 300, 300 Ω) equipped with foam tips. The earphones were driven by a TDT HB7 headphone driver (Tucker-Davis Technologies, Alachua, FL, USA), receiving signals from an RME Fireface UCX soundcard (RME, Haimhausen, Germany). All audio signals were presented at a sampling rate of 44.1 kHz. The experimental setup was acoustically calibrated using a Brüel & Kjær (B&K) Type 4157 Ear Simulator equipped with a DB-2012 Ear Mould Adaptor and a secure cap. The signal was pre-amplified by a B&K Type 2669 Microphone Preamplifier and connected to a B&K Type 2610 Measuring Amplifier. Prior to measurements, the measuring amplifier was calibrated using a B&K Type 4231 Sound Calibrator. The speech-shaped OLSA noise was used to calibrate the behavioral speech audiometry, the MEG OLSA task, as well as the long-term average sound level of the audiobooks. Visual stimuli (fixation cross, response options, and comprehension questions) were projected using a PROPixx MRI/MEG projector (VPIXX, Saint-Bruno-de-Montarville, Canada). Participant responses were recorded using a fiber-optic button box (RESPONSEPixx/MRI, VPIXX) connected to a DATAPixx3 data acquisition system (VPIXX), which also managed the visual output timing. The experiments were implemented using Python 3.8 and PsychoPy (v2024.1.4) (Peirce et al. 2019). To ensure low-latency playback, audio stimuli for all tasks (Behavioral OLSA, MEG OLSA and MEG Audiobooks) were handled by the audio engine SoundMexPro (Berg 2024). Trigger signals were generated simultaneously as an aligned audio channel from the soundcard, transmitted as a digital signal to a custom-made trigger box, and subsequently to the MEG system. This custom-built FPGA-based trigger box utilized the clock of the Fireface UCX to convert the SPDIF output into a 5 V TTL trigger input to the MEG. In the OLSA task, triggers marked the precise onset of each sentence, whereas the audiobook task utilized synchronization triggers at 1 s intervals aligned with the audio stream.

### Data analysis

Data analysis was performed for all 21 subjects, conceptually following Vanthornhout et al. (2018) and is depicted in Figure 1. Subject sub-00 was included as a pilot, and sub-01 and sub-19 were included despite slightly increased thresholds in one ear, as their inclusion did not alter results. For analysis, the clean speech condition was remapped to 0 dB for sub-00, and the *−*40 dB and 0 dB conditions were effectively remapped to 0 % and 100 % intelligibility for all subjects, as behavioral performance was consistently at floor (*≤*1 %) or ceiling (*≥*98 %) at these levels. The main analysis was performed on the pooled data from both sessions (ses-01, ses-02) to increase statistical power and the amount of available data, as preliminary results were comparable across both recording sessions. Therefore, the dataset comprised four audiobook runs (two audiobooks, each split into two parts) yielding approximately 30 minutes of continuous speech per participant, and six OLSA runs totaling 1080 sentences (200 sentences per intelligibility condition, with the exception of the *−*40 dB condition, which contained 80 sentences, except sub-00).

#### Signal processing and preprocessing

Neural recordings were preprocessed using MNE-Python’s (Gramfort et al. 2013) implementation of Maxwell filtering (Taulu and Kajola 2005; Taulu and Simola 2006), which included spatiotemporal signal space separation (tSSS), head movement correction using cHPI data, and transformation to a common head position (relative to the *ses-01 task-olsa run-02* recording). This transformation ensured consistent sensor-to-head alignment across all recordings and sessions, further facilitating coregistration. Subsequent analysis was conducted in MATLAB using the FieldTrip toolbox (Oostenveld et al. 2011). Continuous MEG and EEG signals for both audiobook and OLSA tasks were filtered in the delta band (0.5 Hz to 4 Hz) using a windowed-sinc finite impulse response filter (firws) (Widmann et al. 2015). Following filtering, audiobook data were segmented into continuous 60-second epochs, downsampled to 64 Hz, and concatenated into a single dataset. Conversely, OLSA recordings were filtered using the same parameters, subsequently cut into epochs corresponding to each sentence (0 s to 0.5 s post-stimulus) and downsampled to 64 Hz. Three sensor modalities were defined for comparison: (1) **MEG**, comprising all 306 channels (magnetometers and gradiometers), (2) **EEG**, comprising all 76 channels (standard 64-channel cap plus 12 near-ear electrodes), which were re-referenced to a common average after filtering and (3) **ear-EEG**, containing 16 near-ear channels using electrodes around the ears (8 per ear) as a subset of the full EEG montage. To ensure robustness and avoid reliance on single channels, the ear-EEG data were re-referenced separately for the left and right ears using the average potential of the respective ear’s channels. To maintain methodological simplicity, recorded EOG and ECG data were not utilized for artifact removal, as the linear decoding approach proved robust without these additional corrections. Envelopes from speech stimuli were extracted following Biesmans et al. (2016) using a Gammatone filterbank with 28 channels spaced by one equivalent rectangular bandwidth between 5 Hz and 5000 Hz. The absolute output of each channel was raised to the power of 0.6, averaged across channels to derive a broadband envelope, and downsampled to 1000 Hz. Finally, the audio envelopes were filtered in the delta band (0.5 Hz to 4 Hz) matching the neural data processing and downsampled to 64 Hz for analysis.

#### Decoder analysis

The decoding analysis aimed to predict speech envelopes from multivariate neural recordings using a backward linear model or simply decoder. We utilized the widely-adopted mTRF-Toolbox for MATLAB (Crosse et al. 2016; Crosse et al. 2021) to reconstruct the speech envelope from neural data using integration latencies between 0 ms and 400 ms. Prior to model training, both data modalities were normalized: audio envelopes were scaled to a range of [*−*1, 1] based on their maximum absolute value, while MEG and EEG data were *z*-scored globally rather than on a channel-by-channel basis. Specifically, this involved calculating a single mean and standard deviation across all MEG or EEG channels and trials to ensure that spatial topographies and relative differences between sensors remained intact. Crucially, to preserve these topographic patterns within the MEG data, *z*-score statistics were calculated across all combined magnetometer or gradiometer channels. The decoder was trained and evaluated on audiobook data pooled across both sessions (approximately 32 minutes, divided into 32 trials) for each individual subject. A cross-validation scheme was employed where 80 % of the randomized trials were utilized for training and parameter optimization (regularization parameter *λ*), while the remaining 20 % served as a held-out test set to verify performance under matched versus mismatched audio envelope conditions. Following optimization, the final model was trained on the full training set using the selected regularization parameter. To derive an objective measure of speech intelligibility as suggested by Vanthornhout et al. (2018), the trained decoder was applied to the OLSA dataset. For each SNR condition, valid speech segments (excluding breaks before and after the sentence) were concatenated to form continuous vectors corresponding to 200 sentences (or 80 sentences for the *−*40 dB condition). We then computed the reconstruction accuracy in terms of Spearman correlation between the reconstructed and original envelope. Additionally, three distinct data conditions were used to distinguish true neural tracking from a null distribution and to enable further statistical inference:

1. **Original correlation:** This value represents the primary measure of neural tracking, calculated as the Spearman correlation between the reconstructed and original speech envelopes for the full, ordered sequence of sentences. Each subject is represented by a single reconstruction accuracy value per SNR level.
2. **Bootstrapped correlation:** To estimate the variance of the observed accuracies, a distribution of correlation values was generated for each SNR via bootstrapping. Predicted and original envelopes were sampled sentence-wise with replacement and concatenated to match the original number of sentences per SNR. This process was repeated 1000 times to produce a distribution of reconstruction accuracies.
3. **Null distribution:** To establish a chance-level baseline, an identical bootstrapping procedure was employed (1000 repetitions per SNR) but with an additional temporal randomization step. Each concatenated reconstructed sentence envelope was circularly shifted by at least 1 s relative to the audio envelope. This procedure disrupts the precise time-locking between the prediction and the stimulus while preserving the global statistical properties of the signals.

Representative envelopes, the corresponding reconstructions and the analysis for an example subject are depicted in Figure 2.

**Figure 2:**
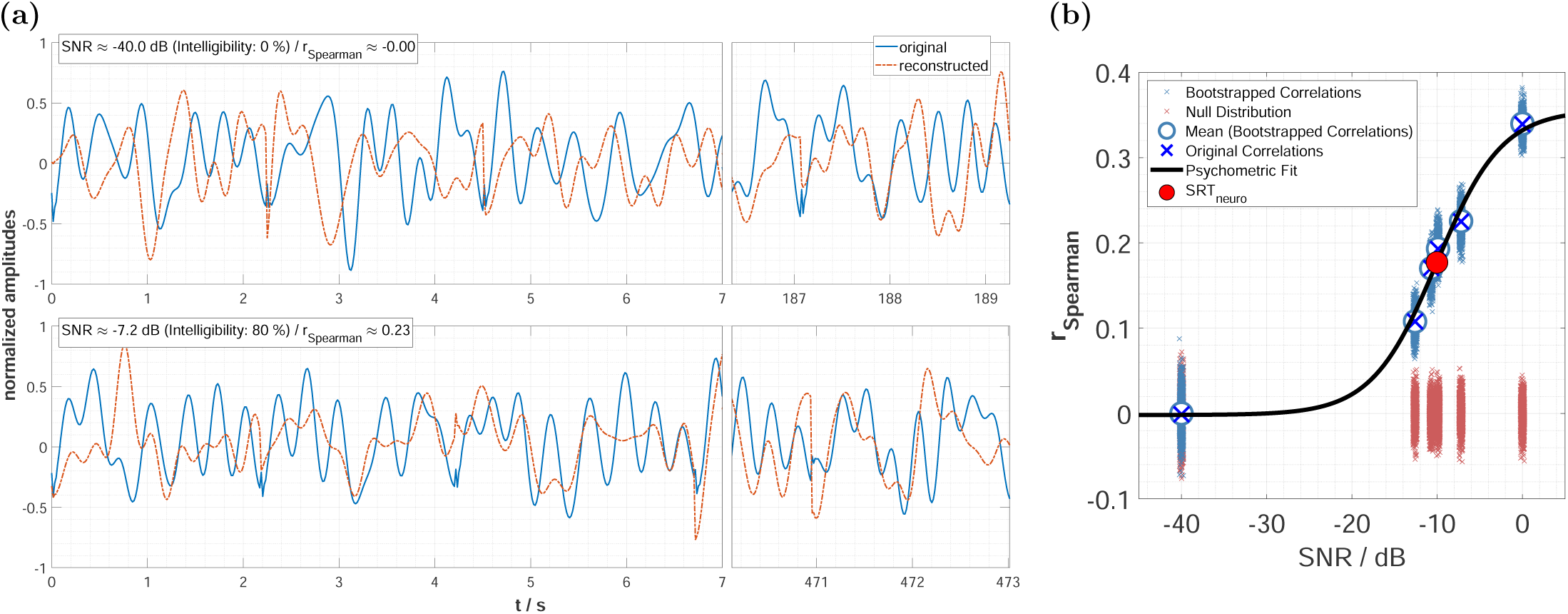
Representative envelope reconstructions and data analysis for sub-10 (MEG, pooled session): **(a)** The original speech envelopes (blue solid) are compared to those reconstructed by the decoder (red dashed). Two exemplary conditions are shown: 0 % intelligibility (upper panel) and 80 % intelligibility (lower panel). Correlations were computed by concatenating segments only containing speech, comprising 80 sentences for the 0 % condition and 200 sentences for all other intelligibility levels. This process results in the sharp transitions visible between segments. Due to the total signal length, only the initial 7 s and final 3 s of the concatenated streams are displayed. **(b)** Correlation values as a function of SNR. The estimated Spearman correlation coefficients derived from the reconstructions in panel (a) are depicted as individual original correlation values (large dark blue crosses). These are shown alongside the bootstrapped correlation distribution (small blue crosses) and its mean (large blue circle). The null distribution is shown in red. A psychometric function (black line) is fitted to original correlation values to estimate the neural speech reception threshold (SRT_neuro_, red circle).

#### Statistical analysis

To account for the skewness of correlation distributions, all reconstruction accuracies (Spearman’s *ρ*) were transformed into *z*-scores using Fisher’s *z*-transformation. For each sensor modality, neural tracking sensitivity was assessed via 21 specific comparisons: 15 pairwise comparisons between the six intelligibility levels and 6 comparisons comparing each level against the null distributions. These comparisons were evaluated at both the group- and subject-levels, with all resulting *p*-values corrected for multiple comparisons using the Holm-Bonferroni procedure. Group-level comparisons were computed using the original correlation values and the means of the individual null distributions. We employed one-sided parametric paired *t*-tests to evaluate directional hypotheses (e.g., higher intelligibility yields greater accuracy). Normality of the difference scores was verified via the Lilliefors test. Effect sizes were quantified using Hedges’ *g_av_* for within-subject designs:

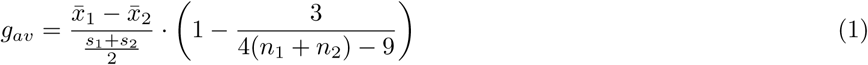

where *x̄*_1_ and *x̄*_2_ denote the grand average means across subjects, *s*_1_ and *s*_2_ represent the standard deviations of the group data, and *n*_1_ and *n*_2_ are the number of subjects per condition. *P*-values were corrected for *n*_tests_ = 63 across the three sensor modalities. Subject-level comparisons utilized individual bootstrapped correlations and null distributions. These were evaluated using one-sided Welch’s *t*-tests to account for unequal variances between the distributions. Effect sizes were computed using Cohen’s *d_s_*for independent samples:

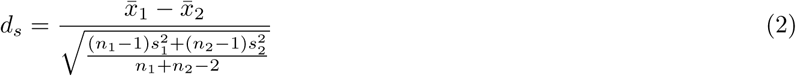

where *x̄*_1_ and *x̄*_2_ are the means of the respective distributions, *s*_1_ and *s*_2_ are the individual standard deviations, and *n*_1_, *n*_2_ denote the number of samples (1000). *P*-values were corrected for *n*_tests_ = 1305 (21 subjects *×* 21 comparisons *×* 3 modalities, sub-00 missing lowest SNR) across all subjects, conditions and modalities. To evaluate relative sensitivity across sensor modalities, linear regressions (intercept constrained to origin) estimated the scaling relationship between reconstruction accuracies for each sensor modality pair. Additionally, Pearson correlations were computed between the original correlation values of all sensor pairings (MEG vs. EEG, MEG vs. ear-EEG, and EEG vs. ear-EEG). Correlations were calculated separately for each of the six intelligibility levels and for all levels pooled together. This resulted in 21 total tests (3 pairings *×* 7 conditions), with *p*-values adjusted using the Holm-Bonferroni procedure.

#### Psychometric function fit

To compare behavioral and neurophysiological thresholds (SRT_beh_ and SRT_neuro_), a sigmoid function was fitted to the original correlation values for each subject. Following Vanthornhout et al. (2018), reconstruction accuracy *S* was modeled as a function of SNR *x*:

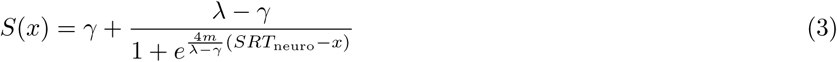

where SRT_neuro_ is the threshold at the inflection point, *m* is the slope, *γ* is the lower asymptote (guess rate), and *λ* is the upper asymptote (maximum accuracy). The fitting procedure, adapted from Borges et al. (2025b) and Borges et al. (2025c), constrained parameters based on individual data: *γ* was restricted to the subject-specific 95 % confidence interval of the null distribution pooled across all SNRs. The upper asymptote *λ* was bounded between the lower bound of the subject-specific 95 % null distribution interval and 105 % of the maximum observed accuracy to account for measurement noise and to mitigate potential underestimation of the ceiling performance. Additionally, SRT_neuro_ was bounded between *−*40 and 10 dB, and the slope *m* was constrained to be non-negative. Optimization was performed using a Nonlinear Least Squares Trust-Region algorithm in MATLAB. To avoid local minima, the procedure was repeated 100 times per subject with the initial SRT_neuro_ set to the group-mean behavioral threshold (*−*9.8 dB). The final parameters were defined as the mean of the 10 fits yielding the highest *R*^2^ values. Test-retest reliability of the estimated SRT_neuro_ was assessed by performing 100 additional fits on resampled data. For each SNR, 10 reconstruction values were drawn randomly from the individual bootstrapped correlation distributions to generate a distribution of threshold estimates and determine their associated standard deviation. Fits were excluded from further analysis if any of the following six criteria were not met. The first three validated the robust neural tracking of the input data: (i) the mean individual effect sizes of the 15 intelligibility pair contrasts were greater than 0.8, (ii) the mean effect size of the 5 non-zero intelligibility levels against the null distribution exceeded 0.8, and (iii) no more than two original correlation values fell outside the range defined by the 100 % intelligibility and the null distribution mean value. The remaining three criteria assessed the validity of the fit itself: (iv) *R*^2^ *>* 0.5, (v) SRT_neuro_ was at least 10 dB from the parameter boundaries (*−*30 dB *< SRT* _neuro_ *<* 0 dB), and (vi) the SRT_neuro_ estimate was not identified as an outlier. Validated SRT_neuro_ and SRT_beh_ estimates were compared using Pearson correlations and by computing the differences, with all *p*-values corrected using False Discovery Rate (FDR) (*n*_tests_ = 3).

#### Monotonicity optimization

Following the approach of Vanthornhout et al. (2018), we optimized the decoder’s frequency bands and integration time windows by maximizing a monotonicity metric, defined as the degree to which the original reconstruction accuracy increases as a function of the SNR. This metric was calculated for each subject and sensor modality as the Spearman correlation coefficient between SNRs and reconstruction accuracies, weighted by the dynamic range of the observed values to account for varying performance ceilings across MEG, EEG, and ear-EEG. Negative correlation results were clamped to 0. This weighting was necessary to resolve non-unique maxima where the Spearman coefficient alone lacked the sensitivity to distinguish between parameter sets with similar trends but different absolute accuracy levels. The optimal parameters were identified via a grid search comprising 10 frequency band combinations derived from lower cutoffs of (0.5, 4, 8, 15) Hz and upper cutoffs of (4, 8, 15, 30) Hz, covering the delta (0.5 Hz to 4 Hz), theta (4 Hz to 8 Hz), alpha (8 Hz to 15 Hz), and beta (15 Hz to 30 Hz) bands. Additionally, we evaluated 10 integration windows using combinations of start offsets (0, 75, 150, 250) ms and end offsets (75, 150, 250, 400) ms. This approach encompassed a comprehensive range of early-to-late neural response latencies while limiting the total number of parameter combinations for the grid search. Optimal parameters for the frequency band and integration window were then applied globally to all subjects.

### Data and Code Availability

The dataset will be made publicly available on the OpenNeuro platform following publication, accompanied by a comprehensive data descriptor. In the interim, the data supporting the findings of this study are available from the corresponding author upon reasonable request. The analysis scripts used in this study are open-source and publicly accessible via GitHub under the BSD 3-Clause License (GitHub repository link will be added later) and archived on Zenodo (Zenodo DOI will be added later) for long-term reproducibility.

Data analysis was performed using a combination of local and remote computing resources. Processing was conducted on a laptop (Lenovo Yoga Pro 9i 16, Windows 11 Home) and a high-performance Linux server (Red Hat Enterprise Linux 9.7) to enable parallel batch processing across participants using 96 logical cores.

## Results

The following results evaluate neural envelope tracking across the three sensor modalities: MEG, EEG, and ear-EEG. We first briefly describe the results of the optimization of the frequency bands and decoder integration windows. Following this, neural tracking is quantified as a function of SNR and speech intelligibility, followed by an assessment of relative sensitivity across the three modalities. We further investigate tracking robustness by analyzing effect sizes and statistical contrasts between intelligibility levels and null distributions at both the group- and subject-levels. Finally, we compare measured behavioral speech reception thresholds with objectively derived neurophysiological thresholds.

### Monotonicity optimization: Frequency band and integration window

The monotonicity optimization for the decoder parameters yielded results consistent with prior reports (Vanthornhout et al. 2018). Across all sensor modalities (MEG, EEG, and ear-EEG), the delta frequency band (0.5 Hz to 4 Hz) achieved the highest monotonicity scores according to our weighted metric. In contrast to the shorter integration windows (e.g., 0 ms to 75 ms) emphasized in previous work, our grid search revealed that extending the integration window to include later latencies was beneficial. Specifically, the full range of 0 ms to 400 ms yielded the highest monotonicity, suggesting that later neural response components contribute to the envelope reconstruction. Consequently, these optimized parameters were utilized for the entirety of the subsequent analysis.

### Neural envelope tracking

#### Neural envelope tracking across SNR and intelligibility

Neural tracking as a function of SNR and speech intelligibility is depicted in Figure 3. In the upper panel (a), reconstruction accuracies consistently increased with SNR for all sensor modalities. Reconstruction accuracies were highest for MEG (reaching *ρ ≈* 0.5 for the best subject), followed by EEG (up to 0.3) and ear-EEG (up to 0.2). Group-level Spearman correlations between SNR and reconstruction accuracy were highly significant (*p <* 0.001) for all modalities (MEG: 0.87, EEG: 0.83, ear-EEG: 0.73). The null distributions yielded accuracies near zero across all SNRs and sensors, confirming the absence of spurious tracking. In the lower panel (b), individual SNRs were mapped onto the six fixed intelligibility levels. For all sensor modalities, reconstruction accuracies increased with intelligibility, with group-level grand averages showing a strictly monotonic increase. Correlations between reconstruction accuracy and intelligibility remained at a high level for all modalities (MEG: 0.87, EEG: 0.78, ear-EEG: 0.73). The variance in reconstruction performance increased at higher intelligibilities, being most pronounced for MEG int the 0 % intelligibility condition. Group-level paired *t*-tests showed that all conditions with intelligibility *≥* 20 % were significantly higher than their respective null distributions, whereas the 0 % condition was statistically indistinguishable from the noise floor. Overall, correlation coefficients were highest for MEG, followed by EEG and ear-EEG, verifying a robust relationship between neural tracking depending on both SNR and speech intelligibility.

**Figure 3:**
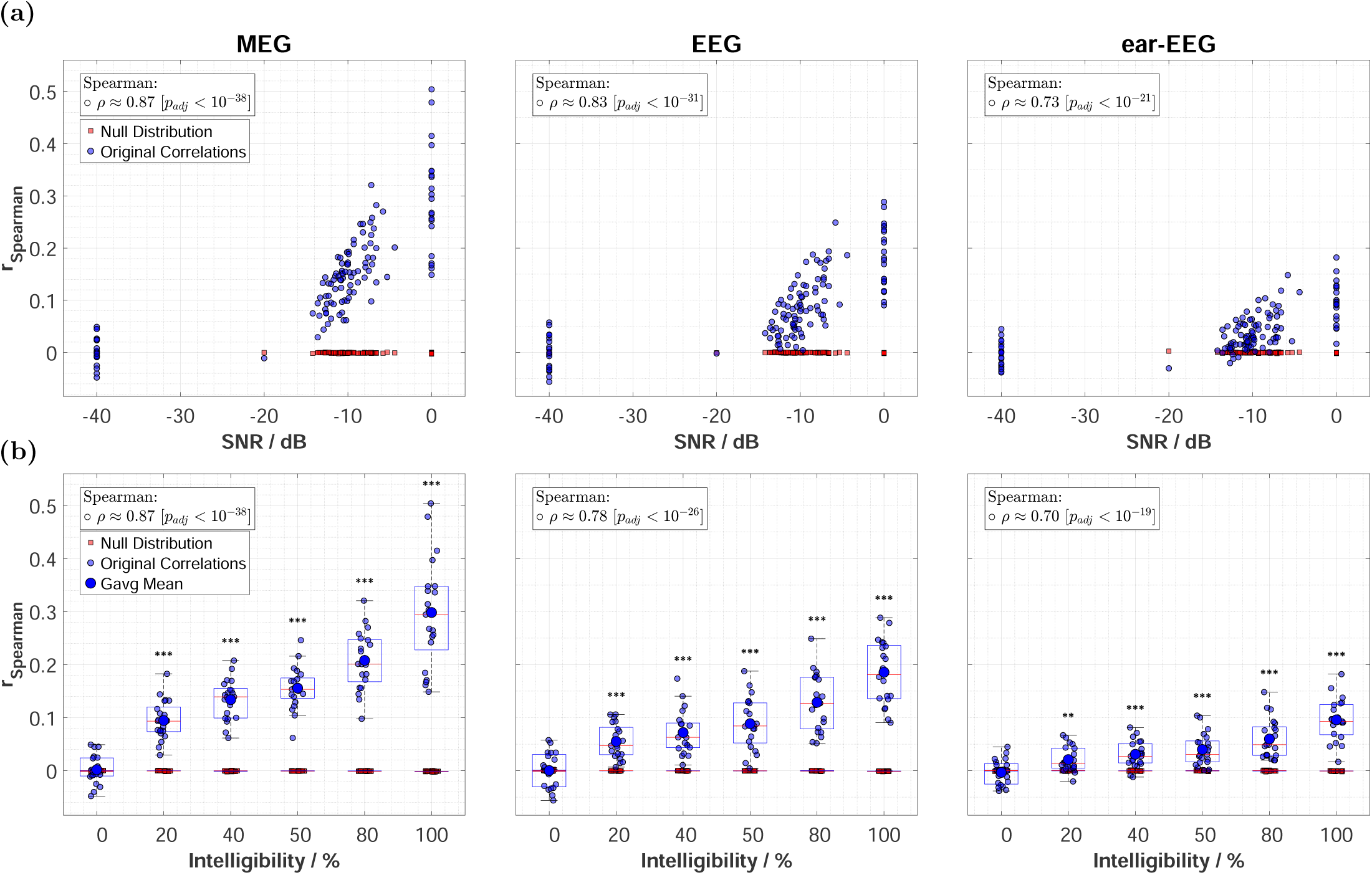
Neural envelope tracking as a function of SNR and speech intelligibility for MEG, EEG, and ear-EEG. Panels (a) and (b) display identical data points for original correlations (blue circles) and individual means of the null distributions (red squares), arranged by individual SNRs and fixed intelligibility levels, respectively. Inset text in both panels displays Spearman correlations (*ρ*) and adjusted *p*-values, which were FDR-corrected (*n*_tests_ = 3). **(a)** Effect of SNR: Reconstruction accuracy is plotted against individual SNRs. **(b)** Effect of intelligibility: data from panel (a) are grouped into the six fixed intelligibility levels. Box plots display the distribution statistics: the central red line indicates the median, dark blue circles represent the grand average mean, box edges indicate the 25th and 75th percentiles, and whiskers extend to the most extreme non-outlier data points. Asterisks indicate the reconstruction accuracies significantly higher than the null distributions based on group-level contrasts (paired *t*-test; *** *p <* 0.001, ** *p <* 0.01, * *p <* 0.05), with *p*-values corrected using the Holm-Bonferroni procedure (*n*_tests_ = 63).

#### Neural envelope tracking consistency between modalities

The consistency of reconstruction accuracies across sensor modalities is illustrated in Figure 4. Pearson correlation coefficients (*ρ*) were calculated and corrected for multiple comparisons for each of the six intelligibility levels individually and for all levels pooled together. Pooled analyses across all intelligibility levels revealed significant positive relationships (*p <* 0.001) for all pairings: MEG–EEG (*ρ* = 0.87), MEG–ear-EEG (*ρ* = 0.78), and EEG–ear-EEG (*ρ* = 0.88). These high coefficients reflect a consistent increase in neural tracking with SNR between all modalities. To quantify relative sensitivity, linear regressions were fitted with the intercept set to zero. The resulting slopes indicate the relative performance ratios between the sensor modalities: MEG achieved approximately 50 % higher reconstruction accuracy than EEG (slope = 1.51), and nearly three times higher accuracy than ear-EEG (slope = 2.72). The full EEG montage provided accuracies roughly twice as large as the ear-EEG subset (slope = 1.81). An analysis within specific intelligibility groups showed that while overall trends remained positive, correlations were not consistently significant at individual levels. This suggests that high neural tracking in one modality does not strictly predict a proportionately high level in another within a single intelligibility condition for a subject, a trend most prominent in pairings involving MEG. In the 0 % (unintelligible) condition, no significant correlations were found, which is consistent with the expected noise-floor behavior. Excluding this condition, significant correlations were observed in three out of five levels for MEG–EEG, but only in one level for MEG–ear-EEG. Notably, all five remaining correlations between ear-EEG and EEG remained significant (*p <* 0.05). This reflects their shared measurement basis, since the ear-EEG sensors are a subset of the full EEG montage.

**Figure 4:**
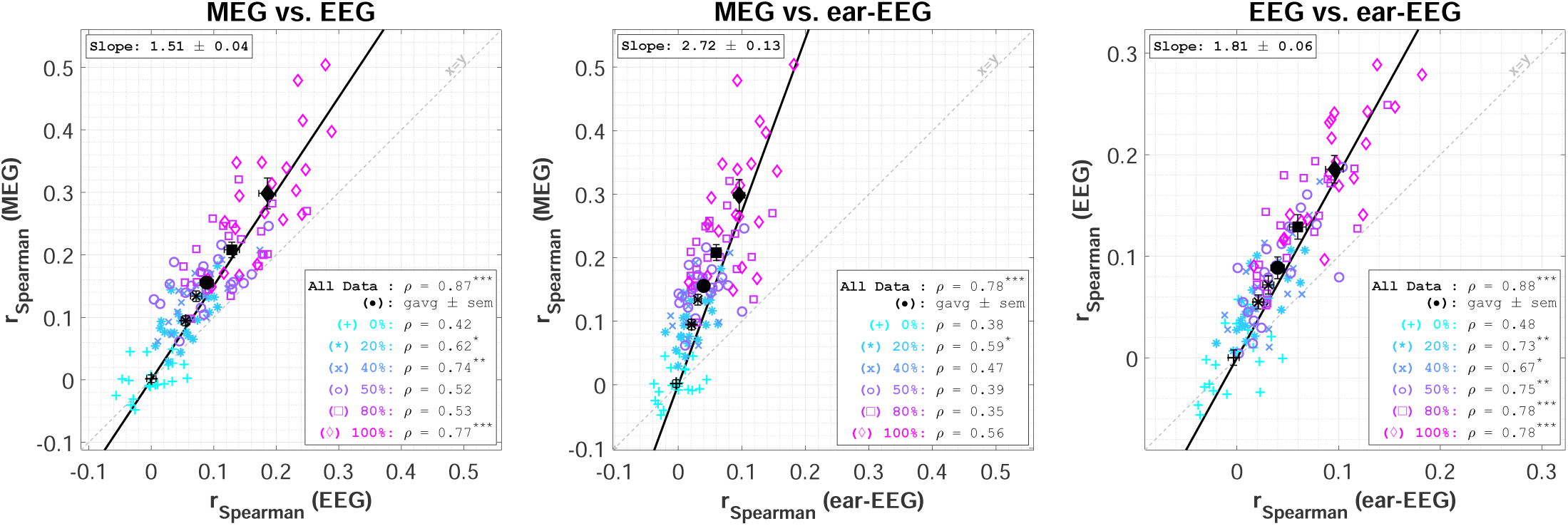
Between-sensor consistency of reconstruction accuracies. Scatter plots compare original correlation values (*r*_Spearman_) across the three sensor modality pairings: MEG vs. EEG, MEG vs. ear-EEG, and EEG vs. ear-EEG. Each marker represents an individual subject at a specific intelligibility level (0%–100%), with colors and marker types indicating the respective levels. Grand averages across all subjects per level (*±* SEM) are shown as bold black markers with error bars. Inset text reports Pearson correlation coefficients (*ρ*) for the pooled dataset (All Data) and individual intelligibility subgroups. Asterisks denote significant correlations after Holm-Bonferroni (*n*_tests_ = 21; *** *p <* 0.001, ** *p <* 0.01, * *p <* 0.05). Solid black lines indicate linear regression fits to the pooled data of all levels (intercept set to origin), with estimated slopes *±* standard error provided in each panel. The identity line (*x* = *y*) is plotted in dashed light gray for reference.

#### Within-modality sensitivities

To further evaluate the sensitivity of each configuration, effect sizes and statistical comparisons between intelligibility levels, including contrasts against null distributions, were computed at the group- and subject-levels. Group-level results are presented in Figure 5. As shown in the heatmaps in Figure 5 (a), nearly all pairwise comparisons showed statistically significant differences for all three modalities. This confirms that MEG, EEG, and ear-EEG are all capable of distinguishing neural tracking across different intelligibility levels. Specifically, significant results along the super-diagonal (contrasts between neighboring intelligibility levels) demonstrate that all modalities can resolve small increments in speech intelligibility at the group level. The comparison between 0 % intelligibility and the null distribution was non-significant across all modalities, confirming expected noise-floor behavior. The left-most columns of the heatmaps, corresponding to Figure 3 (b), show comparisons between the null distribution and each intelligibility level. MEG exhibited the largest effect sizes (Hedges’ *g_av_*), reaching values up to 8, compared to approximately 6 for EEG and 4 for ear-EEG. While all modalities demonstrated high statistical power, MEG achieved *p <* 0.001 for nearly all contrasts. EEG maintained significance across the diagonal, whereas for ear-EEG, the 40 % vs. 50 % contrast did not reach significance. Figure 5 (b) illustrates the consistency of these group-level effect sizes between modality pairings. Effect sizes for all 21 comparisons were significantly correlated across all pairings, with the highest similarity observed between EEG and ear-EEG (*ρ* = 0.97), followed by MEG–EEG (*ρ* = 0.85) and MEG–ear-EEG (*ρ* = 0.77). Linear regression revealed that MEG was approximately 1.4 times more sensitive in terms of effect sizes than EEG and nearly twice as sensitive as ear-EEG. EEG was approximately 1.3 times more sensitive than the ear-EEG configuration.

**Figure 5:**
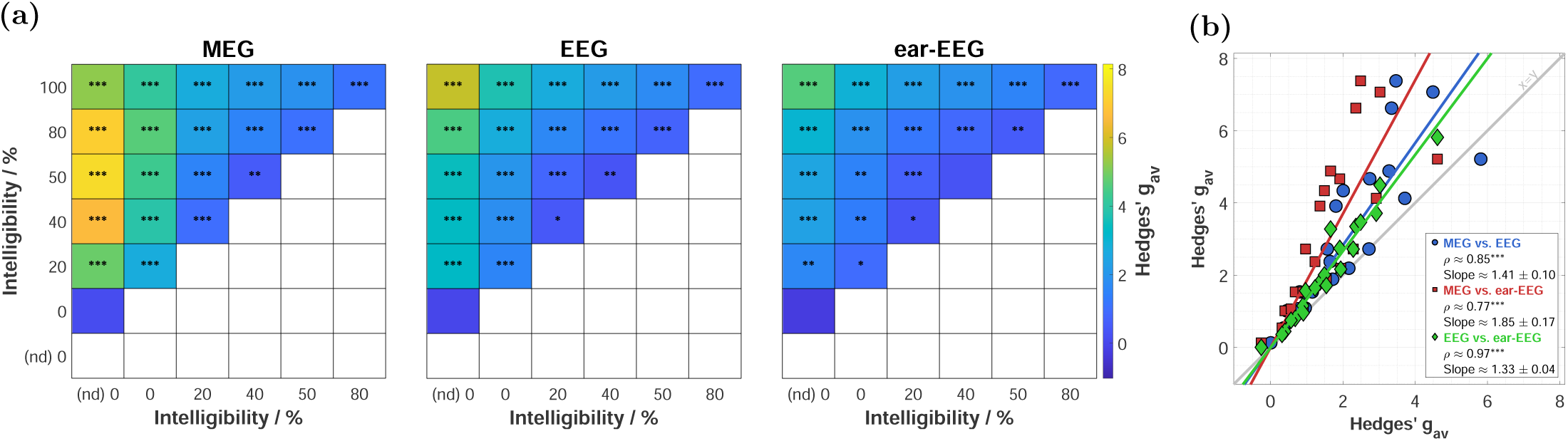
Group-level effect sizes for comparisons between intelligibility levels and against null distributions (nd). **(a)** Heatmaps for MEG, EEG, and ear-EEG. Color intensity represents the effect size (Hedges’ *g_av_*) for all 21 contrasts. Statistical significance of pairwise comparisons, assessed via paired-sample *t*-tests, is indicated by asterisks (* *p <* 0.05, ** *p <* 0.01, *** *p <* 0.001). All *p*-values were Holm-Bonferroni corrected for multiple comparisons (*n*_tests_ = 63). **(b)** Between-sensor consistency of effect sizes: Effect sizes from panel (a) are rearranged into scatter plots for each modality pairing. Inset text displays the Pearson correlation coefficients (*ρ*) and linear regression slopes (intercept set to origin) *±* standard error. *P*-values were adjusted using the FDR procedure (*n*_tests_ = 3).

Subject-level sensitivity is summarized in Figure 6, which displays the number of subjects reaching significant thresholds (*p <* 0.05) for each contrast. MEG demonstrated the highest detection rates and greatest stability, with nearly all comparisons reaching significance for all subjects. For EEG and ear-EEG, detection rates decreased along the super-diagonal (neighboring conditions). For neighbor-condition contrasts, MEG showed significance on average for 19 participants, compared to approximately 17 for EEG and 15 for ear-EEG. Interestingly, for all modalities, the contrast between 0 % intelligibility and the null distribution remained significant for roughly one-third of the subjects, suggesting a detectable statistical difference even at 0 % intelligibility.

**Figure 6:**
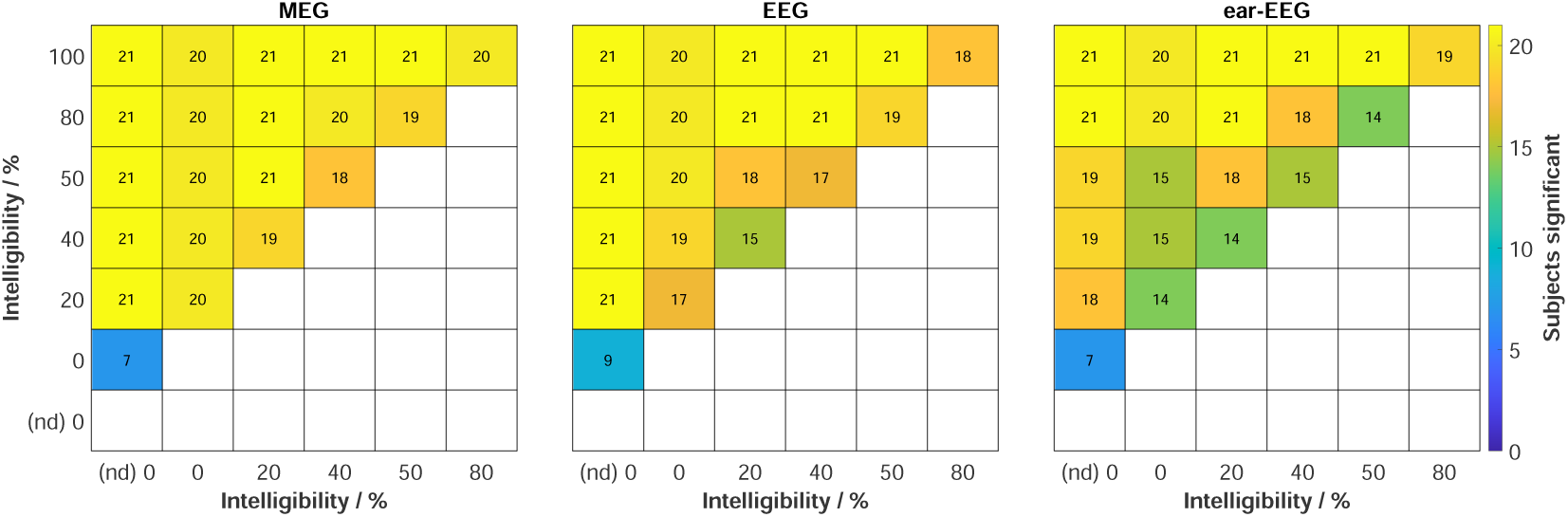
Subject-level counts for significant comparisons between intelligibility levels and against null distributions (nd). Heatmaps display the number of subjects exhibiting significant comparisons for MEG, EEG, and ear-EEG. The first column of each heatmap indicates the number of subjects with bootstrapped correlations significantly higher than their respective null distributions. The remaining cells indicate the number of subjects showing significant pairwise differences in reconstruction accuracy between intelligibility levels. Statistical significance was determined via one-sided Welch’s *t*-tests (*p <* 0.05), with *p*-values corrected for multiple comparisons (*n*_tests_ = 1305) using the Holm-Bonferroni procedure.

### Behavioral vs. neurophysiological threshold estimates

Behavioral thresholds were narrowly distributed between *−*11.6 dB to *−*7.1 dB (spread: 4.5 dB). In contrast, neurophysiological thresholds exhibited wider ranges that expanded as the number of sensors decreased: MEG (*−*12.1 dB to *−*7.0 dB), EEG (*−*12.7 dB to *−*5.5 dB), and ear-EEG (*−*13.2 dB to *−*5.9 dB). Based on the validity criteria, SRT_neuro_ was successfully estimated for 20 subjects in MEG, 21 in EEG, and 18 in ear-EEG. As shown in Figure 7 (a), no significant Pearson correlations were found between SRT_beh_ and SRT_neuro_ for any modality. The combined test-retest reliability corridor reached a maximum of 1.54 dB, accounting for the 1 dB OLSA variance and the modality-specific SRT_neuro_ fit reliability (MEG: 0.32 dB, EEG: 0.53 dB, ear-EEG: 1.1 dB). These values assume independence between threshold measurements with omitted covariance. Most SRT_beh_–SRT_neuro_ pairs fell outside this corridor, indicating a lack of a systematic relationship at the individual level. Correlations were predominantly negative, suggesting a trend where SRT_neuro_ thresholds were too high (i.e., less negative), effectively underestimating the hearing threshold. Despite the lack of behavioral correlation, SRT_neuro_ estimates demonstrated significant correlations between sensor modalities (Figure 7 (b)). Positive correlations were observed for all pairings, with the highest consistency between MEG–EEG (*ρ* = 0.73), followed by MEG–ear-EEG (*ρ* = 0.60) and EEG–ear-EEG (*ρ* = 0.50). The distribution of differences (SRT_beh_ – SRT_neuro_) is depicted in Figure 7 (c) and summarized in Table 2. Mean differences were minimal, ranging from 0.1 dB to a slight *−*0.9 dB offset for ear-EEG. This indicates the absence of systematic bias for MEG and EEG, with only a marginal systematic bias observed for ear-EEG. Absolute differences *≤* 3 dB were observed for 17 subjects in MEG, 18 in EEG and 13 in ear-EEG. Standard deviations of these differences increased from 2.0 dB (MEG) to 2.5 dB (ear-EEG), reflecting the higher uncertainty associated with configurations utilizing fewer sensors.

**Table 2:** Summary performance metrics per sensor modality, including the number of subjects having valid fits, the count of subjects with absolute differences *≤* 3 dB, mean, median, standard deviation (Std), and Root Mean Squared Error (RMSE).

| Metric | MEG | EEG | ear-EEG |
| --- | --- | --- | --- |
| $N_{\text{Valid}}$ (out of 21) | 20 | 21 | 18 |
| $N_{\leq 3\text{dB}}$ (out of 21) | 17 | 18 | 13 |
| Mean / dB | 0.1 | $-0.5$ | $-0.9$ |
| Median / dB | 0.2 | $-0.6$ | $-1.6$ |
| Std / dB | 2.0 | 2.4 | 2.5 |
| RMSE / dB | 2.1 | 2.4 | 2.4 |

**Figure 7:**
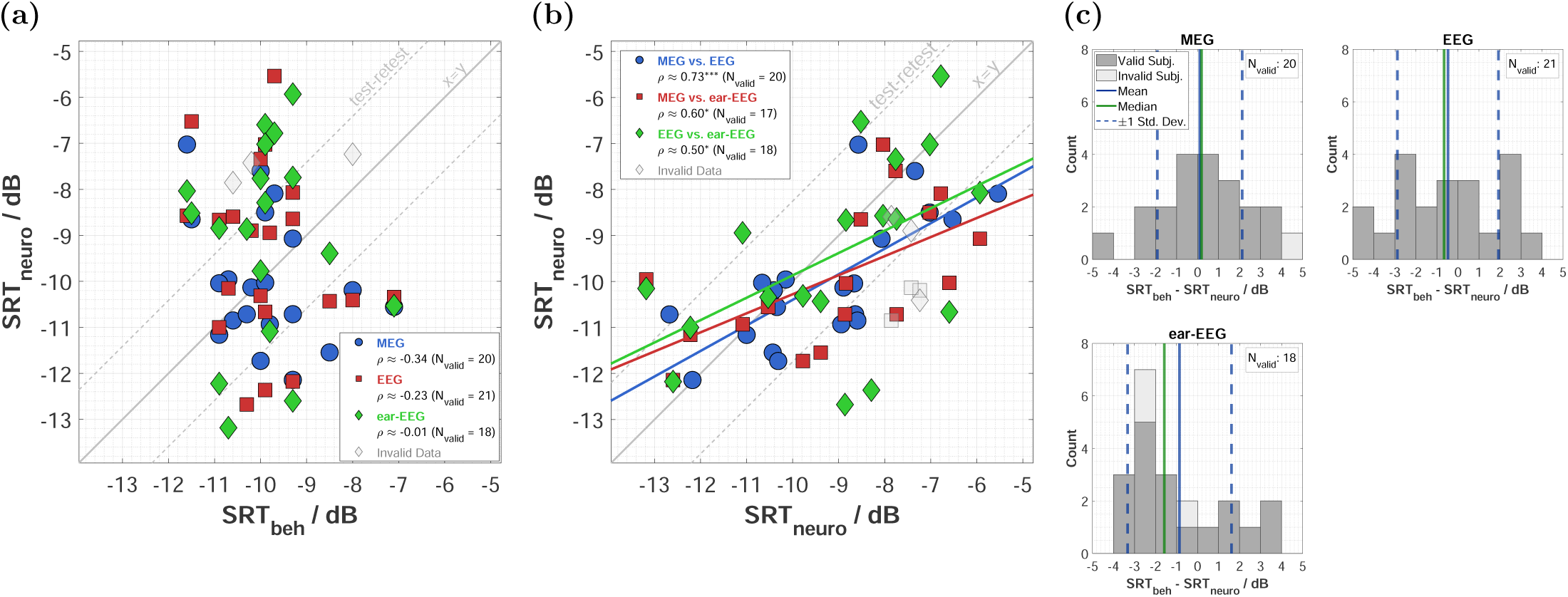
Comparison between behavioral and neurophysiological speech reception thresholds (SRT_beh_ and SRT_neuro_): **(a)** Correlations between SRT_beh_ and SRT_neuro_ for MEG, EEG, and ear-EEG. Each data point represents a single subject, with color and marker coding per sensor modality. Textboxes indicate Pearson correlation coefficients (*ρ*) and the number of valid threshold estimates (*N*_valid_). An identity line (*x* = *y*) is shown with a shaded test-retest reliability corridor. Invalid estimates are indicated in light gray. **(b)** Between-sensor consistency of SRT_neuro_ across sensor modality pairings, including Pearson correlation coefficients and linear regression parameters. Shaded regions denote the reliability corridor relative to the identity line (*x* = *y*). **(c)** Distribution of differences (SRT_beh_ - SRT_neuro_) for each sensor modality. Histograms display the median (green line), mean (solid blue line), and *±*1 standard deviation (dashed blue lines). Invalid threshold estimates are shown in light gray. All *p*-values were corrected for multiple comparisons across the three sensor modalities (*n*_tests_ = 3) using the FDR procedure.

## Discussion

We observed consistent neural tracking, effect sizes, and estimated speech reception thresholds across MEG, EEG, and ear-EEG, demonstrating high similarity between the three methods. Reconstruction accuracies increased consistently alongside SNR and intelligibility. Specifically, MEG yielded reconstruction accuracies approximately 1.5 times higher than EEG and 3 times higher than ear-EEG. All modalities successfully discriminated small SNR increments with high effect sizes. At the individual level, tracking robustness followed the same order, proving most consistent for MEG, followed by EEG and ear-EEG. Neurophysiological thresholds (SRT_neuro_) were successfully derived for nearly all participants as unbiased estimators of behavioral performance. These threshold estimates remained within a 5 dB margin of behavioral values for all modalities. However, ear-EEG exhibited a slight bias and reduced reliability. No significant correlation was found between purely envelope-based SRT_neuro_ and individual SRT_beh_ for any modality.

### Neural envelope tracking

Across all three modalities, we observed a clear increase in neural tracking in terms of higher reconstruction accuracies with increasing SNR and speech intelligibility. This concurrent increase is expected, as individual SNRs were anchored to fixed intelligibility levels via a monotonically increasing psychometric function.

This enhancement of neural tracking, particularly in the delta and theta bands, is well-established in the literature (for a systematic review, see Ratelle and Tremblay (2025)). Prior studies have consistently reported this effect using linear encoder or decoder models, cross-correlation functions, coherence, and phase-locking values. Scalp-EEG is the most widely used modality for these linear models (Das et al. 2018; Decruy et al. 2020a; Vanthornhout et al. 2019; Lesenfants et al. 2019; Vanthornhout et al. 2018; Verschueren et al. 2020; Wang et al. 2020; Zhang et al. 2023; Zou et al. 2019; Decruy et al. 2020b; Decruy et al. 2019; Borges et al. 2025b; Borges et al. 2025a; Van Hirtum et al. 2023), followed by MEG (Ding and Simon 2013; Karunathilake et al. 2023a; Presacco et al. 2016; Presacco et al. 2019), and more recently, ear-EEG (Borges et al. 2025c; Borges et al. 2025a).

### Reconstruction accuracies and effect sizes

Our results follow a clear sensitivity hierarchy (MEG *>* cap-EEG *>* ear-EEG) aligning well with the measured magnitudes and inter-subject variability reported in existing literature. Specifically, Destoky et al. (2019) demonstrated a 50 % sensitivity advantage for MEG over EEG in noiseless conditions. We observed a nearly identical advantage in our noisy conditions, as indicated by the slope of a linear fit between modalities across SNRs. To quantify the separability between SNR levels, we evaluated effect sizes, which followed the same hierarchy. While group-level separability was evident across all modalities (with the exception of the 40 % and 50 % contrast in ear-EEG), ear-EEG exhibited a roughly 33 % reduction in effect size compared to cap-EEG. This finding is consistent with the 21 % to 44 % signal loss reported for auditory event-related potentials when comparing ear-EEG to cap-EEG (Meiser and Bleichner 2022), confirming that the reduced spatial coverage of around-ear montages results in a decrease in tracking sensitivity. However, the high correlations in reconstruction accuracies and effect sizes across all three sensors suggest they capture shared underlying neural information.

### Individual robustness

Beyond group-level significance, individual analyses demonstrate the consistency of each modality. MEG proved the most robust modality, resolving neural tracking curves, defined as significant differences between adjacent intelligibility levels, across participants. The hierarchy reflects the varying spatial coverage and sensitivity of the modalities. The linear decoder may particularly benefit from MEG’s high sensor density and signal richness. While both MEG and EEG provide full-head coverage, magnetic fields remain undistorted by the skull and varying tissue conductivities. Therefore, each channel provides more unique, independent information with significantly less spatial smearing than EEG (Goldenholz et al. 2009; Baillet 2017). Furthermore, the MEG system utilizes significantly more sensors than the EEG-cap (306 vs. 76). In contrast, ear-EEG is restricted to fewer channels within the temporal regions. Despite this spatial limitation, its proximity to the temporal cortex, a region central to processing acoustic features like the speech envelope (Brodbeck et al. 2018a; Kubanek et al. 2013), ensures high local sensitivity (Meiser et al. 2020). The observation that ear-EEG resolved significant neural tracking in 75 % of individuals underscores its viability for detecting subtle SNR increments and potential. However, this restricted coverage reduces sensitivity to neural sources and is more susceptible to individual anatomical differences resulting increased inter-subject variability (Meiser et al. 2020; Meiser and Bleichner 2022; Meiser et al. 2024). While studies suggest that performance in AAD can be maintained or even improved with fewer EEG channels (Mirkovic et al. 2015; Montoya-Martínez et al. 2021), augmenting ear-EEG with even a few scalp electrodes can boost performance by over 10 % (Geirnaert et al. 2025), highlighting the clear advantage of broader spatial sampling. To prioritize real-world applicability, we utilized 16 around-ear channels, referencing each ear independently against its respective local average. This configuration ensures galvanic separation, modeling the decoupled nature of bilateral hearing aids and increasing the setup’s relevance for real-world wearable applications. An exhaustive evaluation of alternative ear-EEG montages and referencing schemes was beyond the scope of the present study.

### Objective speech audiometry

While most research into objective SRT prediction has focused on scalp-EEG, fewer studies have explored ear-EEG, and to our knowledge, none have utilized MEG. Following similar studies (Lesenfants et al. 2019; Muncke et al. 2022; Borges et al. 2025b; Borges et al. 2025c; Deoisres et al. 2025), we evaluated the psychometric fits and the distribution of SRT differences for each sensor modality.

Valid SRT_neuro_ estimates were obtained for nearly all participants (MEG: 20/21, EEG: 21/21), mirroring the robust neural tracking observed across modalities. The lower validity for ear-EEG (18/21) aligns with its reduced sensitivity and with reports that standalone around-ear montages slightly underperform relative to full-cap or combined in-ear and around-ear configurations (Borges et al. 2025c). Precision was moderate to high, with the majority of MEG (17/21) and EEG (18/21) estimates falling within a 3 dB margin of behavioral thresholds. Although ear-EEG robustness was lower (13/21), all participants remained within a 5 dB corridor. This level of precision exceeds that of non-attentive paradigms (Deoisres et al. 2025) or grand-average encoder models (Lesenfants et al. 2019), though it trails optimized ear-EEG configurations reported in the literature (Borges et al. 2025c). Mean and median differences near 0 dB suggest that SRT_neuro_ serves as an unbiased estimator for MEG and EEG. Ear-EEG exhibited a minor bias (*−*1.6 dB), slightly exceeding the best-case median bias (*−*0.15 dB) reported in the literature (Borges et al. 2025c). In contrast to previous reports (Vanthornhout et al. 2018; Accou et al. 2021), we found no significant correlation between individual SRT_beh_ and SRT_neuro_ in any modality.

### Methodological considerations and design validation

As summarized in Table 3, our experimental design aligns with existing literature regarding SRT_beh_, experimental design and model configurations.

**Table 3:** Comparison of related studies. Only configurations most closely matching the current setup are included for studies reporting multiple parameters.

| Study | $SRT_{beh}$ / dB (Range) | SNR Selection (Range) | Model (Frequency Band, Integration Window) | Data (Train / Test) |
| --- | --- | --- | --- | --- |
| Vanthornhout et al. (2018)* | $-7.4 \pm 1.3$ (−9.9 to −4.7) | Fixed: 3–5–7 SNRs (−12.5 to −1 dB) including clean speech | Decoder (Delta, 0 ms to 75 ms) | Comparable to Lesenfants et al. (2019) |
| Lesenfants et al. (2019)* | $-7.1 \pm 1.5$ (−10.3 to −4.7) | Fixed: 5–7 SNRs including clean speech (−12.5 to 2.5 dB) | Encoder (Delta, 0 ms to 400 ms) | 14 min AB / 120–160 matrix sent. (6 min to 8 min) per SNR |
| Borges et al. (2025b) | −5.4 (−6.0 to −3.3) | Individual: $SRT_{beh} \pm$ (0 dB, 2 dB, 4 dB) and clean speech | Decoder (Delta+Theta, −100 ms to 350 ms) | 16 min AB / 16 min AB per SNR |
| Deoisres et al. (2025) | $-10.8 \pm 1.1$ (−12.3 to −9.3) | Fixed: −15 dB, −10 dB, −5 dB and 0 dB and clean speech | Decoder (Delta, 0 ms to 300 ms) | 15 min AB / 8 min AB per SNR |
| Current Study | $-9.8 \pm 1.1$ (−11.6 to −7.1) | Individual: $SRT_{beh} +$ (−2.7 dB, −0.8 dB, 0 dB, 2.7 dB), Fixed: −40 dB and 0 dB | Decoder (Delta, 0 ms to 400 ms) | 30 min AB / 80–200 matrix sent. per SNR (sessions combined) |
| *Studies utilized the same underlying dataset. AB: Audiobook; sent.: Sentences. |  |  |  |  |

#### Behavioral thresholds

Our behavioral thresholds and their distribution align with typical normal-hearing cohorts reported for the OLSA (Nuesse et al. 2019; Deoisres et al. 2025). The slightly lower thresholds relative to Nuesse et al. (2019) (*−*8.6 dB) are likely attributable to diotic rather than monaural presentation, as earphone choice plays a negligible role (Arlinger and Billermark 1997). Our results reflect higher sensitivity than the open-set test thresholds reported by Borges et al. (2025b), where lower variability (2.7 dB *<* 4.5 dB) may have contributed to smaller deviations between SRT_beh_ and SRT_neuro_.

#### Experimental design

To characterize neural tracking, we utilized six SNR levels: four individualized SNRs anchored to SRT_beh_ to sample the slope, supplemented by two fixed anchors at *−*40 dB and 0 dB to capture floor and ceiling effects. This sampling, in particular comparable to the 2 dB steps in Borges et al. (2025b), demonstrates that envelope tracking can resolve SNR increments as small as 1 dB across all modalities. Additional analysis revealed that single-session (15 min training) yielded results similar to the pooled 30 min data, suggesting that further data increases had minimal impact. Moreover, as more sentences were concatenated, reconstruction accuracy remained stable while variability decreased, consistent with observations in AAD (Lopez-Gordo et al. 2025). This indicates that shorter testing durations per SNR may suffice for reliable neural tracking estimates. Differences in performance may also stem from using non-homogeneous training and test materials (audiobooks vs. matrix sentences). While matched materials benefited Borges et al. (2025b), a similar advantage was not observed by Deoisres et al. (2025). Regarding the setup, simultaneous acquisition required participants to wear an EEG-cap within the MEG. While this configuration limits movement more than the standalone EEG recordings cited in Table 3, we aimed to mitigate potential impacts on signal quality by prioritizing participant comfort.

#### Model parameters

To maximize reconstruction accuracy, we employed individual linear backward models integrating data across all available channels and optimized time windows. This engineering-driven approach prioritizes robust envelope reconstruction over the neurophysiological interpretability of forward-model Temporal Response Functions (TRFs) (Haufe et al. 2014). Specifically, our delta-band decoder utilized a 0 ms to 400 ms integration window to capture both early and late neural components, extending the 0 ms to 75 ms window used by Vanthornhout et al. (2018) to align with other reported models. This configuration yielded the most robust and monotonic reconstruction accuracies across our dataset. While combined delta—theta analysis performed similarly to the delta band alone (Borges et al. 2025b), exclusive theta-band tracking proved inferior, compromising threshold validity and monotonicity (Deoisres et al. 2025). These results underscore the delta band’s essential role in robust neural envelope tracking.

#### Importance of asymptotes

Unlike behavioral recognition naturally bounded between 0 and 100 %, neural reconstruction limits are mathematically defined by correlation coefficients within the range of *−*1 to 1. This makes them more intricate to define. Subject specific boundaries must be carefully established because the asymptotes significantly impacts SRT_neuro_ by shifting the function’s range. This effect is magnified as psychometric slopes become shallower. We utilized fixed SNRs to characterize these limits. The lower asymptote had a negligible impact. At the group level, reconstruction at *−*40 dB did not differ from the null distribution. Although one third of participants showed significant differences between these conditions, the absolute values were too small to meaningfully alter the fitted range and threshold estimates. Importantly, inter-subject variability in reconstruction accuracy scaled with SNR and became most pronounced at the highest signal levels, a trend particularly evident in the MEG data. Such variability is critical because these high SNR levels define the upper asymptote. Unlike studies that used clean speech as a reference, we used the 0 dB condition as a proxy because it provided 100 % intelligibility at approximately 10 dB above the mean SRT_beh_. While some literature suggests that clean speech may yield higher reconstruction accuracies than 0 dB (Decruy et al. 2019; Karunathilake et al. 2023a), other evidence indicates accuracy saturation or only minor deviations between clean and high SNR speech (Herrmann 2025a; Herrmann 2025b; Ding and Simon 2013; Van Hirtum et al. 2023). To mitigate potential underestimation, the fitted ceiling was allowed to reach 105 % of the maximal recorded accuracy. Notably, audiobook data proved unsuitable as a ceiling for matrix sentences due to inherently higher reconstruction accuracies (Verschueren et al. 2020). We recommend including an additional clean speech reference using matched materials for future recordings (Borges et al. 2025b).

#### Artifact correction

Artifact correction was omitted to streamline the analysis pipeline because its benefit for continuous speech tracking remains debatable. While Holtze et al. (2022) observed no improvement in classification accuracy, Geirnaert et al. (2025) found that artifact removal specifically benefited ear-EEG. Conversely, cap-EEG showed negligible gains in reconstruction accuracy or threshold prediction (Borges et al. 2025c). We evaluated this impact by applying Independent Component Analysis (ICA) to Maxwell filtered MEG data and performing both bad-channel interpolation and ICA on the full EEG montage. For the ear-EEG subset, this served as an upper performance bound approximation by utilizing information from all EEG channels. This analysis yielded only minor differences for MEG and cap-EEG. For ear-EEG, artifact correction increased the number of valid threshold estimates (+3) and overall reliability, but it failed to establish a significant correlation between SRT_beh_ and SRT_neuro_ or improve accuracy within the 3 dB margin. Crucially, a control decoder trained on EOG and ECG channels failed to reconstruct the speech envelope across all SNRs. This lack of predictive power confirms that systematic artifacts did not drive the observed neural tracking, further justifying the use of uncorrected signals.

### Impact of attention

Attention significantly modulates reconstruction accuracy (Vanthornhout et al. 2019; Reetzke et al. 2021). Although neural tracking persists during passive listening, accuracies remain lower than in active conditions which is commonly exploited in AAD (O’sullivan et al. 2015; Ding and Simon 2012; Golumbic et al. 2013). Diminished attention can lower the upper asymptote and distort SRT_neuro_ estimates. This likely contributes to the reduced accuracy and higher variability observed in non-attentive paradigms (Deoisres et al. 2025). Following previous work (Vanthornhout et al. 2018; Borges et al. 2025b), we monitored engagement through content related questions. We used jittered question intervals and frequent intercom contact to maintain alertness in the MEG scanner. However, participants still reported fatigue during repetitive OLSA blocks. Using audiobooks with varying SNRs may provide a more engaging and ecologically valid alternative (Borges et al. 2025b).

### Audibility vs. intelligibility

Whether neural envelope tracking reflects speech comprehension remains a subject of debate. Although it correlates with behavioral measures, it appears to be a necessary but insufficient condition for intelligibility (Gillis et al. 2022). Because intelligibility typically covaries with acoustic properties, distinguishing higher-level speech processing from the tracking of physical stimulus characteristics remains challenging. Studies manipulating intelligibility through vocoding and prior exposure while holding acoustics constant often find no tracking differences between conditions (Kösem et al. 2023; Karunathilake et al. 2023b; Millman et al. 2015). Robust tracking of non-speech signals (Deoisres et al. 2025) and foreign languages (Etard and Reichenbach 2019; Gillis et al. 2023) further suggests the envelope primarily reflects acoustic processing. Conversely, Iotzov and Parra (2019) reported increased tracking during congruent versus incongruent audio visual speech despite identical acoustics.

### Future perspectives

To isolate meaningful speech processing, features such as phoneme or word surprisal, cohort entropy and word frequency are employed to capture neural representations beyond acoustics. However, analyses must strictly control for acoustic confounds to maintain interpretability (Gillis et al. 2021; Gillis et al. 2023). Beyond psychometric fits, alternative methods for estimating SRT_neuro_ do exist. For instance, Lesenfants et al. (2019) utilized theta-band zero-crossings by combining phonetic features with spectrograms to minimize deviations from behavioral thresholds. Other strategies, such as analyzing changes in TRF morphology across SNRs, have successfully placed the majority of participants within a 2 dB corridor of their behavioral SRT (Muncke et al. 2022). Furthermore, deep learning architectures, such as convolutional neural networks, have demonstrated high generalizability and predictive accuracy (Accou et al. 2021; Na et al. 2022). By mapping brain signals to these richer linguistic features, advanced models can capture the non-linearities of brain dynamics, potentially overcoming the inherent limitations of linear approaches in future applications (Puffay et al. 2023a; Puffay et al. 2023b; Puffay et al. 2024). While linear envelope-based decoders demonstrate broad validity across age groups (Borges et al. 2025a), our evidence suggests they may be insufficient for precise subject-level diagnostics (MacIntyre et al. 2024; Panela et al. 2024), even when utilizing the superior spatial sampling of MEG. Ultimately, leveraging the spatial and temporal precision of MEG to capture speech-related brain dynamics using advanced features may facilitate the refinement of these complex models, thereby providing a framework for clinical translation through more portable EEG solutions.

## Conclusion

This study evaluated neural envelope tracking and its potential for objective speech reception threshold (SRT_neuro_) estimation across MEG, scalp-EEG and ear-EEG. Robust tracking, scaling consistently with SNR and speech intelligibility, was observed in all modalities. While MEG and scalp-EEG yielded the highest signal quality and reliability, ear-EEG proved to be a viable alternative for capturing stable neural responses. Although SRT_neuro_ estimates fell within a 5 dB margin of behavioral SRT values, no significant correlation was found between the two for any modality. This suggests that linear modeling of the acoustic envelope may be insufficient as a standalone predictor for individual behavioral thresholds, as it is limited by high inter-subject variability. To enhance diagnostic precision, future research should integrate higher-level speech features and advanced modeling, leveraging MEG’s spatial resolution to better isolate specific neural contributions to speech processing.

## Acknowledgments

This work was supported by the Neuroimaging Unit of the Carl von Ossietzky Universität Oldenburg, funded by grants from the German Research Foundation (3T MRI INST 184/152-1 FUGG and MEG INST 184/148-1 FUGG). The authors would like to thank all participants for their patience and endurance. We also acknowledge Dilek Erdil and Ole Hausendorf for their help with data acquisition, Andreas Spiegler for technical support with the experimental setup, and Daniel Berg for assisting with the Python implementation of the OLSA framework. Furthermore, we thank Rieke Ollermann and Hongmei Hu for their insightful feedback and discussions. During the preparation of this manuscript, the authors used Gemini 3.1 Pro to improve the language and readability of selected sentences. After using this AI tool, the authors reviewed and edited the text as needed and take full responsibility for the final content of the publication.

## Author contributions

CRediT: Till Habersetzer: Conceptualization, Methodology, Software, Validation, Formal analysis, Investigation, Data Curation, Writing - Original Draft, Writing - Review & Editing, Visualization, Project administration. Andreas Radeloff: Conceptualization, Writing - Review & Editing, Funding acquisition. Bernd T. Meyer: Writing - Review & Editing, Project administration, Funding acquisition.

## Funding

This research is funded by Deutsche Forschungsgemeinschaft (DFG, German Research Foundation) under Germany’s Excellence Strategy - EXC 2177/2 - Project ID 390895286 and by Forschungspoolmittel Potentialbereich mHealth from the School VI, Medicine and Health Sciences at the Carl von Ossietzky Universität Oldenburg (PB mHealth 2020-13).

## Notes

### Competing Interest Statement

The authors have declared no competing interest.

## References

Accou, B., M. Jalilpour Monesi, H. Van Hamme, and T. Francart (2021). “Predicting speech intelligibility from EEG in a non-linear classification paradigm”. In: Journal of Neural Engineering 18.6, p. 066008.

Arlinger, S. D. and E. Billermark (1997). “Hearing thresholds for speech using insert earphones versus supra-aural earphones”. In: Scandinavian audiology 26.3, pp. 151–154.

Baillet, S. (2017). “Magnetoencephalography for brain electrophysiology and imaging”. In: Nature neuroscience 20.3, pp. 327–339.

Berg, D. (Sept. 2024). SoundMexPro. Version 3.1.0.0. doi: 10.5281/zenodo.13847645. url: https://doi.org/10.5281/zenodo.13847645.

Biesmans, W., N. Das, T. Francart, and A. Bertrand (2016). “Auditory-inspired speech envelope extraction methods for improved EEG-based auditory attention detection in a cocktail party scenario”. In: IEEE Trans. Neural Syst. Rehabil. Eng. 25.5, pp. 402–412.

Borges, H. B., E. Alickovic, C. B. Christensen, P. Kidmose, and J. Zaar (2025a). “Age-Related Differences in EEG-Based Speech Reception Threshold Estimation Using Scalp and Ear-EEG”. In: Trends in hearing 29, p. 23312165251372462.

Borges, H. B., J. Zaar, E. Alickovic, C. B. Christensen, and P. Kidmose (2025b). “Speech reception threshold estimation via EEG-based continuous speech envelope reconstruction”. In: European Journal of Neuroscience 61.6, e70083.

Borges, H. B., J. Zaar, E. Alickovic, C. B. Christensen, and P. Kidmose (2025c). “The speech reception threshold can be estimated using EEG electrodes in and around the ear”. In: Journal of Neural Engineering 22.5, p. 056008.

Brand, T. and B. Kollmeier (2002). “Efficient adaptive procedures for threshold and concurrent slope estimates for psychophysics and speech intelligibility tests”. In: The Journal of the Acoustical Society of America 111.6, pp. 2801– 2810.

Brodbeck, C., L. E. Hong, and J. Z. Simon (2018a). “Rapid transformation from auditory to linguistic representations of continuous speech”. In: Current Biology 28.24, pp. 3976–3983.

Brodbeck, C., A. Presacco, and J. Z. Simon (2018b). “Neural source dynamics of brain responses to continuous stimuli: Speech processing from acoustics to comprehension”. In: NeuroImage 172, pp. 162–174.

Brodbeck, C. and J. Z. Simon (2020). “Continuous speech processing”. In: Current Opinion in Physiology 18, pp. 25–31.

Crosse, M. J., G. M. Di Liberto, A. Bednar, and E. C. Lalor (2016). “The multivariate temporal response function (mTRF) toolbox: a MATLAB toolbox for relating neural signals to continuous stimuli”. In: Frontiers in human neuroscience 10, p. 604.

Crosse, M. J., N. J. Zuk, G. M. Di Liberto, A. R. Nidiffer, S. Molholm, and E. C. Lalor (2021). “Linear modeling of neurophysiological responses to speech and other continuous stimuli: methodological considerations for applied research”. In: Frontiers in neuroscience 15, p. 705621.

Das, N., A. Bertrand, and T. Francart (2018). “EEG-based auditory attention detection: boundary conditions for background noise and speaker positions”. In: Journal of neural engineering 15.6, p. 066017.

Debener, S., R. Emkes, M. De Vos, and M. Bleichner (2015). “Unobtrusive ambulatory EEG using a smartphone and flexible printed electrodes around the ear”. In: Scientific reports 5.1, p. 16743.

Debener, S., F. Minow, R. Emkes, K. Gandras, and M. De Vos (2012). “How about taking a low-cost, small, and wireless EEG for a walk?” In: Psychophysiology 49.11, pp. 1617–1621.

Decruy, L., D. Lesenfants, J. Vanthornhout, and T. Francart (2020a). “Top-down modulation of neural envelope tracking: the interplay with behavioral, self-report and neural measures of listening effort”. In: European Journal of Neuroscience 52.5, pp. 3375–3393.

Decruy, L., J. Vanthornhout, and T. Francart (2019). “Evidence for enhanced neural tracking of the speech envelope underlying age-related speech-in-noise difficulties”. In: Journal of neurophysiology 122.2, pp. 601–615.

Decruy, L., J. Vanthornhout, and T. Francart (2020b). “Hearing impairment is associated with enhanced neural tracking of the speech envelope”. In: Hearing Research 393, p. 107961.

Deoisres, S., G. S. Aljarboa, S. L. Bell, and D. M. Simpson (2025). “Comparing approaches for predicting behavioural speech-in-noise performance using cortical responses to unattended stimuli”. In: Hearing Research 457, p. 109197.

Destoky, F., M. Philippe, J. Bertels, M. Verhasselt, N. Coquelet, M. Vander Ghinst, V. Wens, X. De Tiège, and M. Bourguignon (2019). “Comparing the potential of MEG and EEG to uncover brain tracking of speech temporal envelope”. In: Neuroimage 184, pp. 201–213.

Ding, N. and J. Z. Simon (2012). “Emergence of neural encoding of auditory objects while listening to competing speakers”. In: Proceedings of the National Academy of Sciences 109.29, pp. 11854–11859.

Ding, N. and J. Z. Simon (2013). “Adaptive temporal encoding leads to a background-insensitive cortical representation of speech”. In: Journal of Neuroscience 33.13, pp. 5728–5735.

Etard, O. and T. Reichenbach (2019). “Neural speech tracking in the theta and in the delta frequency band differentially encode clarity and comprehension of speech in noise”. In: Journal of Neuroscience 39.29, pp. 5750–5759.

Geirnaert, S., S. L. Kappel, and P. Kidmose (2025). “A direct comparison of simultaneously recorded scalp, around-ear and in-ear EEG for neural selective auditory attention decoding to speech”. In: Scientific Reports 15.1, p. 41655.

Geirnaert, S., S. Vandecappelle, E. Alickovic, A. De Cheveigne, E. Lalor, B. T. Meyer, S. Miran, T. Francart, and A. Bertrand (2021). “Electroencephalography-based auditory attention decoding: Toward neurosteered hearing devices”. In: IEEE Signal Processing Magazine 38.4, pp. 89–102.

Gillis, M., J. Van Canneyt, T. Francart, and J. Vanthornhout (2022). “Neural tracking as a diagnostic tool to assess the auditory pathway”. In: Hearing Research 426, p. 108607.

Gillis, M., J. Vanthornhout, and T. Francart (2023). “Heard or understood? Neural tracking of language features in a comprehensible story, an incomprehensible story and a word list”. In: eneuro 10.7.

Gillis, M., J. Vanthornhout, J. Z. Simon, T. Francart, and C. Brodbeck (2021). “Neural markers of speech comprehension: measuring EEG tracking of linguistic speech representations, controlling the speech acoustics”. In: Journal of Neuroscience 41.50, pp. 10316–10329.

Goldenholz, D. M., S. P. Ahlfors, M. S. Hämäläinen, D. Sharon, M. Ishitobi, L. M. Vaina, and S. M. Stufflebeam (2009). “Mapping the signal-to-noise-ratios of cortical sources in magnetoencephalography and electroencephalography”. In: Human brain mapping 30.4, pp. 1077–1086.

Golumbic, E. M. Z., N. Ding, S. Bickel, P. Lakatos, C. A. Schevon, G. M. McKhann, R. R. Goodman, R. Emerson, A. D. Mehta, J. Z. Simon, et al. (2013). “Mechanisms underlying selective neuronal tracking of attended speech at a “cocktail party””. In: Neuron 77.5, pp. 980–991.

Gramfort, A., M. Luessi, E. Larson, D. A. Engemann, D. Strohmeier, C. Brodbeck, R. Goj, M. Jas, T. Brooks, L. Parkkonen, et al. (2013). “MEG and EEG data analysis with MNE-Python”. In: Frontiers in Neuroinformatics 7, p. 267.

Gross, J. (2019). “Magnetoencephalography in cognitive neuroscience: a primer”. In: Neuron 104.2, pp. 189–204.

Haufe, S., F. Meinecke, K. Görgen, S. Dähne, J.-D. Haynes, B. Blankertz, and F. Bießmann (2014). “On the interpretation of weight vectors of linear models in multivariate neuroimaging”. In: Neuroimage 87, pp. 96–110.

Herrmann, B. (2025a). “Age-related differences in the impact of background noise on neural speech tracking”. In: Neurobiology of Aging 153, pp. 10–20.

Herrmann, B. (2025b). “Enhanced neural speech tracking through noise indicates stochastic resonance in humans”. In: Elife 13, RP100830.

Holtze, B., M. Rosenkranz, M. Jaeger, S. Debener, and B. Mirkovic (2022). “Ear-EEG measures of auditory attention to continuous speech”. In: Frontiers in Neuroscience 16, p. 869426.

Humes, L. E. (2019). “The World Health Organization’s hearing-impairment grading system: an evaluation for unaided communication in age-related hearing loss”. In: International journal of audiology 58.1, pp. 12–20.

Iotzov, I. and L. C. Parra (2019). “EEG can predict speech intelligibility”. In: Journal of Neural Engineering 16.3, p. 036008.

Karunathilake, I. D., J. L. Dunlap, J. Perera, A. Presacco, L. Decruy, S. Anderson, S. E. Kuchinsky, and J. Z. Simon (2023a). “Effects of aging on cortical representations of continuous speech”. In: Journal of neurophysiology 129.6, pp. 1359–1377.

Karunathilake, I. D., J. P. Kulasingham, and J. Z. Simon (2023b). “Neural tracking measures of speech intelligibility: Manipulating intelligibility while keeping acoustics unchanged”. In: Proceedings of the National Academy of Sciences 120.49, e2309166120.

Kösem, A., B. Dai, J. M. McQueen, and P. Hagoort (2023). “Neural tracking of speech envelope does not unequivocally reflect intelligibility”. In: NeuroImage 272, p. 120040.

Kubanek, J., P. Brunner, A. Gunduz, D. Poeppel, and G. Schalk (2013). “The tracking of speech envelope in the human cortex”. In: PloS one 8.1, e53398.

Lesenfants, D., J. Vanthornhout, E. Verschueren, L. Decruy, and T. Francart (2019). “Predicting individual speech intelligibility from the cortical tracking of acoustic-and phonetic-level speech representations”. In: Hearing research 380, pp. 1–9.

Looney, D., P. Kidmose, C. Park, M. Ungstrup, M. L. Rank, K. Rosenkranz, and D. P. Mandic (2012). “The in-the-ear recording concept: User-centered and wearable brain monitoring”. In: IEEE pulse 3.6, pp. 32–42.

Lopez-Gordo, M. A., S. Geirnaert, and A. Bertrand (2025). “Unsupervised accuracy estimation for brain-computer interfaces based on selective auditory attention decoding”. In: IEEE Transactions on Biomedical Engineering.

Luo, H. and D. Poeppel (2007). “Phase patterns of neuronal responses reliably discriminate speech in human auditory cortex”. In: Neuron 54.6, pp. 1001–1010.

MacIntyre, A. D., R. P. Carlyon, and T. Goehring (2024). “Neural decoding of the speech envelope: effects of intelligibility and spectral degradation”. In: Trends in Hearing 28, p. 23312165241266316.

Malmivuo, J. (2012). “Comparison of the properties of EEG and MEG in detecting the electric activity of the brain”. In: Brain topography 25.1, pp. 1–19.

Meiser, A. and M. G. Bleichner (2022). “Ear-EEG compares well to cap-EEG in recording auditory ERPs: A quantification of signal loss”. In: Journal of Neural Engineering 19.2, p. 026042.

Meiser, A., A. Lena Knoll, and M. G. Bleichner (2024). “High-density ear-EEG for understanding ear-centered EEG”. In: Journal of Neural Engineering 21.1, p. 016001.

Meiser, A., F. Tadel, S. Debener, and M. G. Bleichner (2020). “The sensitivity of ear-EEG: evaluating the source-sensor relationship using forward modeling”. In: Brain topography 33.6, pp. 665–676.

Millman, R. E., S. R. Johnson, and G. Prendergast (2015). “The role of phase-locking to the temporal envelope of speech in auditory perception and speech intelligibility”. In: Journal of cognitive neuroscience 27.3, pp. 533–545.

Mirkovic, B., S. Debener, M. Jaeger, and M. De Vos (2015). “Decoding the attended speech stream with multi-channel EEG: implications for online, daily-life applications”. In: Journal of neural engineering 12.4, p. 046007.

Montoya-Martínez, J., J. Vanthornhout, A. Bertrand, and T. Francart (2021). “Effect of number and placement of EEG electrodes on measurement of neural tracking of speech”. In: Plos one 16.2, e0246769.

Muncke, J., I. Kuruvila, and U. Hoppe (2022). “Prediction of speech intelligibility by means of EEG responses to sentences in noise”. In: Frontiers in Neuroscience 16, p. 876421.

Na, Y., H. Joo, L. T. Trang, L. D. A. Quan, and J. Woo (2022). “Objective speech intelligibility prediction using a deep learning model with continuous speech-evoked cortical auditory responses”. In: Frontiers in Neuroscience 16, p. 906616.

Nicolas-Alonso, L. F. and J. Gomez-Gil (2012). “Brain computer interfaces, a review”. In: sensors 12.2, pp. 1211–1279.

Nuesse, T., B. Wiercinski, T. Brand, and I. Holube (2019). “Measuring speech recognition with a matrix test using synthetic speech”. In: Trends in Hearing 23, p. 2331216519862982.

O’sullivan, J. A., A. J. Power, N. Mesgarani, S. Rajaram, J. J. Foxe, B. G. Shinn-Cunningham, M. Slaney, S. A. Shamma, and E. C. Lalor (2015). “Attentional selection in a cocktail party environment can be decoded from single-trial EEG”. In: Cerebral cortex 25.7, pp. 1697–1706.

Oostenveld, R., P. Fries, E. Maris, and J.-M. Schoffelen (2011). “FieldTrip: open source software for advanced analysis of MEG, EEG, and invasive electrophysiological data”. In: Computational intelligence and neuroscience 2011.1, p. 156869.

Panela, R. A., F. Copelli, and B. Herrmann (2024). “Reliability and generalizability of neural speech tracking in younger and older adults”. In: Neurobiology of Aging 134, pp. 165–180.

Pasley, B. N., S. V. David, N. Mesgarani, A. Flinker, S. A. Shamma, N. E. Crone, R. T. Knight, and E. F. Chang (2012). “Reconstructing speech from human auditory cortex”. In: PLoS biology 10.1, e1001251.

Peelle, J. E. and M. H. Davis (2012). “Neural oscillations carry speech rhythm through to comprehension”. In: Frontiers in psychology 3, p. 320.

Peirce, J., J. R. Gray, S. Simpson, M. MacAskill, R. Höchenberger, H. Sogo, E. Kastman, and J. K. Lindeløv (2019). “PsychoPy2: Experiments in behavior made easy”. In: Behavior research methods 51.1, pp. 195–203.

Presacco, A., J. Z. Simon, and S. Anderson (2016). “Evidence of degraded representation of speech in noise, in the aging midbrain and cortex”. In: Journal of neurophysiology 116.5, pp. 2346–2355.

Presacco, A., J. Z. Simon, and S. Anderson (2019). “Speech-in-noise representation in the aging midbrain and cortex: Effects of hearing loss”. In: PloS one 14.3, e0213899.

Puffay, C., B. Accou, L. Bollens, M. J. Monesi, J. Vanthornhout, H. Van Hamme, and T. Francart (2023a). “Relating EEG to continuous speech using deep neural networks: a review”. In: Journal of Neural Engineering 20.4, p. 041003.

Puffay, C., J. Vanthornhout, M. Gillis, B. Accou, H. Van Hamme, and T. Francart (2023b). “Robust neural tracking of linguistic speech representations using a convolutional neural network”. In: Journal of Neural Engineering 20.4, p. 046040.

Puffay, C., J. Vanthornhout, M. Gillis, P. D. Clercq, B. Accou, H. V. hamme, and T. Francart (2024). “Classifying coherent versus nonsense speech perception from EEG using linguistic speech features”. In: Scientific Reports 14.1, p. 18922.

Ratelle, D. and P. Tremblay (2025). “Neural tracking of continuous speech in adverse acoustic conditions among healthy adults with normal hearing and hearing loss: A systematic review”. In: Hearing Research, p. 109367.

Reetzke, R., G. N. Gnanateja, and B. Chandrasekaran (2021). “Neural tracking of the speech envelope is differentially modulated by attention and language experience”. In: Brain and Language 213, p. 104891.

Silva, F. L. da (2013). “EEG and MEG: relevance to neuroscience”. In: Neuron 80.5, pp. 1112–1128.

Taulu, S. and M. Kajola (2005). “Presentation of electromagnetic multichannel data: the signal space separation method”. In: Journal of Applied Physics 97.12.

Taulu, S. and J. Simola (2006). “Spatiotemporal signal space separation method for rejecting nearby interference in MEG measurements”. In: Physics in Medicine & Biology 51.7, p. 1759.

Van Hirtum, T., B. Somers, E. Verschueren, B. Dieudonné, and T. Francart (2023). “Delta-band neural envelope tracking predicts speech intelligibility in noise in preschoolers”. In: Hearing Research 434, p. 108785.

Vanthornhout, J., L. Decruy, and T. Francart (2019). “Effect of task and attention on neural tracking of speech”. In: Frontiers in neuroscience 13, p. 977.

Vanthornhout, J., L. Decruy, J. Wouters, J. Z. Simon, and T. Francart (2018). “Speech intelligibility predicted from neural entrainment of the speech envelope”. In: Journal of the Association for Research in Otolaryngology 19.2, pp. 181–191.

Verschueren, E., J. Vanthornhout, and T. Francart (2020). “The effect of stimulus choice on an EEG-based objective measure of speech intelligibility”. In: Ear and hearing 41.6, pp. 1586–1597.

Wagener, K. and T. Brand (2005). “Sentence intelligibility in noise for listeners with normal hearing and hearing impairment: Influence of measurement procedure and masking parameters”. In: International journal of audiology 44.3, pp. 144–156.

Wagener, K., T. Brand, and B. Kollmeier (1999). “Development and evaluation of a German sentence test Part III: Evaluation of the Oldenburg sentence test”. In: Zeitschrift Fur Audiologie 38, pp. 86–95.

Wang, L., E. X. Wu, and F. Chen (2020). “Robust EEG-based decoding of auditory attention with high-rms-level speech segments in noisy conditions”. In: Frontiers in human neuroscience 14, p. 557534.

WHO, et al. (2021). World report on hearing. World Health Organization.

Widmann, A., E. Schröger, and B. Maess (2015). “Digital filter design for electrophysiological data–a practical approach”. In: Journal of neuroscience methods 250, pp. 34–46.

Wöstmann, M., L. Fiedler, and J. Obleser (2017). “Tracking the signal, cracking the code: Speech and speech comprehension in non-invasive human electrophysiology”. In: Language, Cognition and Neuroscience 32.7, pp. 855–869.

Zhang, X., J. Li, Z. Li, B. Hong, T. Diao, X. Ma, G. Nolte, A. K. Engel, and D. Zhang (2023). “Leading and following: Noise differently affects semantic and acoustic processing during naturalistic speech comprehension”. In: NeuroImage 282, p. 120404.

Zou, J., J. Feng, T. Xu, P. Jin, C. Luo, J. Zhang, X. Pan, F. Chen, J. Zheng, and N. Ding (2019). “Auditory and language contributions to neural encoding of speech features in noisy environments”. In: NeuroImage 192, pp. 66–75.

